# MEST inhibits ciliary sphingomyelin synthesis to promote tendon stem/progenitor cells osteochondrogenesis in traumatic heterotopic ossification

**DOI:** 10.1101/2025.03.01.640937

**Authors:** Bowen Lai, Yuan Gao, Jianquan Zhao, Zhilong Shen, Rui Gao, Heng Jiang, Xuhui Zhou

**Affiliations:** Department of Orthopedics, Changzheng Hospital, Naval Medical University (Second Military Medical University); Shanghai, 200003, China; Department of Orthopedic Oncology, Changzheng Hospital, Naval Medical University (Second Military Medical University); Shanghai, 200003, China; Translational Research Centre of Orthopedics, Shanghai General Hospital, Shanghai Jiao Tong University School of Medicine; Shanghai, 200080, China

**Keywords:** heterotopic ossification, primary cilia, tendon stem/progenitor cells, mesoderm-specific transcript, sphingomyelin

## Abstract

Traumatic heterotopic ossification (tHO) is a musculoskeletal disorder characterized by ectopic bone formation in soft tissues following trauma. However, the etiology and mechanisms of tHO remain unknown, and no treatments are highly effective for tHO. Here, by performing single-cell RNA sequencing (scRNA-seq) on two tHO mouse models, we found that ciliary Hedgehog (Hh) and PI3K-Akt signaling in tendon stem/progenitor cells (TSPCs) is upregulated during tHO development, leading to the activation of GLI family zinc finger 2 (GLI2) transcription factor, which promotes mesoderm-specific transcript (MEST) gene expression. Untargeted lipidomics using liquid chromatography-tandem mass spectrometry (LC-MS/MS) on detached cilia indicated that MEST could reduce ciliary sphingomyelin (SM) levels by inhibiting SM synthesis in TSPCs, creating a positive feedback loop that amplified ciliary Hh signaling, enhancing TSPCs osteochondrogenesis and driving tHO formation. Targeting ciliary genes intraflagellar transport 88 (IFT88) or ADP ribosylation factor like GTPase 3 (ARL3) restored normal TSPCs osteochondrogenesis in vitro and attenuated ectopic bone formation in vivo by suppressing Hh and PI3K-Akt signaling. AAV-mediated MEST inhibition or exogenous SM administration in vivo also alleviates tHO progression. These findings may provide novel insights into tHO pathogenesis and potential therapeutic strategies.

## INTRODUCTION

Traumatic heterotopic ossification (tHO) is a common musculoskeletal disorder characterized by the formation of ectopic bone in soft tissues, affecting a significant proportion of patients who experience severe injuries or orthopedic procedures^1^. Although most cases of tHO present with localized symptoms such as pain, swelling, and restricted joint movement, severe cases can lead to significant functional impairments, including joint ankylosis and chronic disability, necessitating intensive surgical interventions like amputation^2^. However, the etiology and underlying biological mechanisms of tHO remain unknown. Notably, there are no effective drugs available that can specifically eliminate ectopic bone formation or decelerate the progression of tHO. Therefore, it is necessary to conduct more studies exploring the molecular mechanisms of tHO to provide new strategies for clinical treatment that alleviate the suffering of patients and the burden on medical resources.

Primary cilia are solitary, non-motile microtubule-based organelles that protrude from the cell surface, acting as essential sensory antennae for cells to receive and transduce extracellular signals^3^. They play crucial roles in various developmental and physiological processes, including embryonic patterning, tissue homeostasis, and cellular differentiation^4^. Recent studies have increasingly highlighted the involvement of primary cilia in skeletal biology. For instance, primary cilia in osteocyte function as critical mechanosensors to maintain normal bone formation and help to treat osteoporosis^5^. Our previous study found that primary cilia can positively regulate the process of osteochondral differentiation of stem cells from patients with fibrodysplasia ossificans progressiva (FOP), a genetic form of HO caused by gene point mutations^6^. However, there is currently no research involving the function and regulatory mechanisms of primary cilia in the pathogenesis of tHO.

The recruitment of stem/progenitor cells and their differentiation into osteoblasts or chondrocytes are key steps in the formation of tHO^7^. Tendon stem/progenitor cells (TSPCs) are located between the long and parallel collagen fiber chains of tendon and are surrounded by the extracellular matrix^8^. TSPCs can differentiate into tenocyte tissue (tendon) and non-tenocyte tissues (such as bone, cartilage, and adipose tissue) in different circumstances. Studies have reported that the increased ectopic bone volume in Achilles tendon of rats is accompanied with increased bone morphogenetic protein (BMP) expression and enhanced osteogenesis in TSPCs^9^. Moreover, the inhibition of the mTOR signaling by rapamycin treatment can suppress the osteochondral differentiation of TSPCs and tHO^10^. Thus, it is important to explore whether primary cilia regulate the abnormal osteochondrogenic differentiation in TSPCs and how primary cilia contribute to the progression of tHO via specific mechanisms.

High-throughput sequencing technologies have been used to find potential mechanisms of various diseases in recent years. In our study, we indicated that MEST might be a key downstream molecule in the regulation of tHO by primary cilia by performing single-cell RNA sequencing (scRNA-seq) in two tHO mouse models. MEST (mesoderm-specific transcript) is an imprinted gene predominantly expressed from the paternal allele. The protein encoded by it is widely expressed in mesodermal tissues (e.g., sclerotome-derived structures such as vertebrae, ribs, and cartilage), belongs to the α/β hydrolase superfamily, and has enzymatic activity. A study showed that knocking down MEST in human periodontal ligament stem cells will inhibit the expression of their stem cell markers and proliferation ability and weaken their multi-lineage differentiation (including osteogenesis and chondrogenesis) abilities^11^. MEST can also upregulate mouse mesenchymal stem cells chondrogenesis along with miR-335-5p^12^. However, no study focuses on whether MEST is involved in regulating TSPCs osteogenic or chondrogenic differentiation. Therefore, it is necessary to explore the role of MEST in the development of tHO and the possibility of targeting MEST as a novel therapeutic strategy.

There is a diffusion barrier in the transition zone at the base of the primary cilia, which can restrict the free movement of proteins and lipids in and out of the cilia, resulting in differences in the composition of proteins and lipids on the ciliary membrane and the cytoplasmic membrane. Under the stimulation of mechanical stress or chemical signals, the selective movement of specific lipids in and out of the cilia can dynamically influence the function of receptors on the ciliary surface and the activity of downstream signaling, thereby regulating cell proliferation and differentiation^13^. The disruption of the homeostasis of ciliary lipids has been reported to be associated with a variety of diseases, including Joubert syndrome^14^, Meckel-Gruber syndrome^15^, nephronophthisis^16^, and polycystic kidney disease^17^. Some studies have reported that abnormalities in genes encoding key enzymes involved in cholesterol metabolism can lead to defects in the formation of primary cilia in osteoblasts and in the differentiation of osteoblasts, resulting in craniofacial malformation^18,19^. Since MEST has been found to possess the characteristics of lipase, which can promote the hypertrophy of adipocytes and expand the volume of adipose tissue^20^, MEST may be involved in lipid metabolism in tHO. Exploring the role of MEST in regulating TSPCs ciliary lipids may be of great significance for clarifying the pathogenesis of tHO.

In this study, we used single-cell RNA sequencing (scRNA-seq) on two tHO mouse models:

1) burn/tenotomy in C57BL/6 mice; 2) cardiotoxin (CTX)-induced injury of tibial muscle in *Nse-Bmp4*mice (overexpresses bone morphogenetic protein 4 (BMP4) under the control of neuron-specific enolase (Nse) promoter). We demonstrated that the activity of ciliary Hh and PI3K-Akt signaling of TSPCs is significantly upregulated during tHO development, leading to the activation and nuclear translocation of the transcription factor GLI2, which promotes the expression of the MEST gene. The upregulation of MEST expression reduces ciliary sphingomyelin (SM) abundance by inhibiting SM synthesis, which in turn amplifies the activation of the ciliary Hh signaling. This forms a positive feedback loop that significantly enhances the osteochondrogenesis of TSPCs, thereby driving local ectopic bone deposition and contributing to the development and progression of tHO. Moreover, knocking down ciliary gene or suppressing Hh/PI3K-Akt signaling restored normal TSPCs differentiation. Inhibiting MEST expression or exogenous SM administration in vivo also had great therapeutic effects on tHO. Unraveling these mechanisms may enhance our understanding of the pathogenesis of tHO and pave the way for the development of innovative and effective therapeutic approaches.

## RESULTS

### 1. Primary cilia in stem/progenitor cells contribute to tHO progression

To induce tHO by burn/tenotomy^21^, 6- to 8-week-old male C57BL/6 mice were subjected to a burn injury using a 60°C heated aluminum block on the dorsal skin, followed by resection of the right Achilles tendon and subsequent experiments (Figure 1a). Tendon tissues from the injured sites were harvested at 0 days (prior to injury), and at 3, 7, 21, and 56 days after injury for subsequent histological examination (Figure 1b). H&E staining demonstrated disorganized and irregular tendon fibrous tissue at 3 to 7 days post-injury. Chondrogenic structures were detected at 21 days, and osteogenic structures were observed at 56 days. Masson trichrome staining showed blue-stained tendon collagen fibers, with the progression of structural disorganization and tHO being similar to that observed with HE staining. Von Kossa staining revealed the appearance of brown calcified nodules at 21 days, which became deeply stained by 56 days, suggesting calcium deposition due to new bone formation.

**Figure 1.**
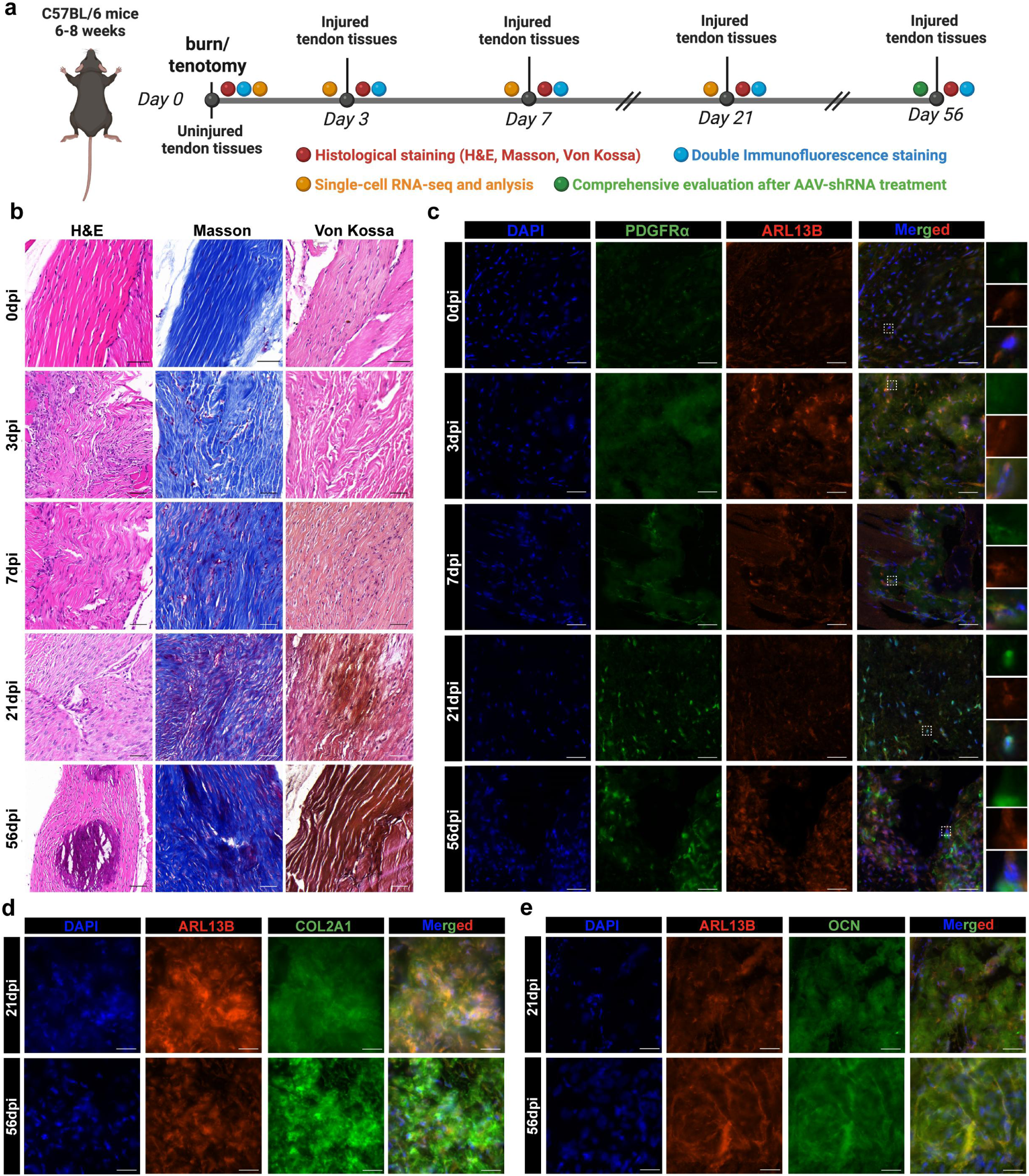
The number of primary cilia-positive cells increases during tHO formation. (a) Schematic diagram of the experimental workflow. (b) H&E, Masson trichrome, and Von Kossa staining of tHO tendon tissues collected from different time points after injury (scale bar 50 µm). (c) Immunofluorescence images of PDGFRα (green) and ARL13B (red) at different time points during tHO development. DAPI staining indicates nuclei (blue), scale bar 30 µm. (d) Immunofluorescence images of COL2A1 (green) and ARL13B (red) in injured sites at 21 and 56 days after tHO. DAPI staining indicates nuclei (blue), scale bar 30 µm. (d) Immunofluorescence images of OCN (green) and ARL13B (red) in injured sites at 21 and 56 days after tHO. DAPI staining indicates nuclei (blue), scale bar 30 µm.

To explore the function of cilia in tHO, we performed immunofluorescence staining on tendon tissues collected from mice at 0, 3, 7, 21, and 56 days post-injury (Figure 1c, S1a). ARL13B, a specific marker for primary cilia, was used to quantify ciliated cells. The percentage of ARL13B-positive cells was 27.21% in the control group (0 days post-injury). After injury, the proportion of ARL13B-positive cells increased to 40.14% at 3 days, 38.07% at 7 days, and 44.42% at 21 days, before declining to 39.10% at 56 days (Figure S1b). These data indicated that ciliated cells increased rapidly during tHO and may contribute to its development. To further identify the potential specific cell types targeted by cilia in the process of tHO, double immunofluorescence staining was performed on tendon tissue samples collected at various time points post-injury (Figure 1c, S1a). The following marker pairs were used: ARL13B with CD31 (endothelial cells marker), ARL13B with F4/80 (macrophages marker), ARL13B with PDGFRα (stem/progenitor cells marker), ARL13B with TNMD (tendon cells marker), and ARL13B with α-SMA (smooth muscle cells marker). Immunofluorescence analysis revealed significant co-localization between ARL13B and all five cell markers, with PDGFRα showing particularly strong association (Figure 1c). Quantitative analysis demonstrated that the proportion of ARL13B+PDGFRα+ double-positive cells was the highest among the five marker combinations (Figure S1c). Specifically, the percentage of ARL13B+PDGFRα+ cells was 55.91% in the control group, and this proportion significantly increased to 78.62% at 3 days, 85.56% at 7 days, 90.17% at 21 days, and 95.49% at 56 days after tHO injury (Figure S1b). We also found that ARL13B is co-stained with both chondrogenic marker gene COL2A1 and osteogenic marker gene OCN at 21 days and 56 days during tHO when osteochondrogenesis have occurred (Figure 1d, e). These data suggested that primary cilia may play a crucial role in tHO development, potentially through their localization in stem/progenitor cells.

Single-cell RNA sequencing (scRNA-seq) can distinguish differences among individual cells in a high-throughput manner, thereby clarifying cellular heterogeneity and underlying molecular mechanisms during disease progression. Therefore, in our study, we reanalyzed a scRNA-seq dataset of injured sites from burn/tenotomy mouse tHO model (GSE126060). The dataset included 12 samples, with three samples per group at four time points: pre-injury (Day 0) and post-injury (Day 3, Day 7, and Day 21). The UMAP plot showed a total of 21,880 cells detected and divided into nine subclusters (Figure 2a) according to the previous study^22^. Bubble plot revealed the marker genes of each cluster, and markers for the Mesenchymal cluster were Pdgfra and Prrx1 (Figure 2b, c). Group-stratified UMAP plots revealed that, as tHO progresses, the percentage of Macrophage cluster cells peaked at 3 days (74.43%) post-injury, later rapidly reduced to 37.10% at 7 days and 15.02% at 21 days post-injury, indicating the evolution of the local inflammatory phase (Figure 2d). In contrast, the Mesenchymal cluster cells exhibited a larger proportion at 7 (43.44%) and 21 days (49.43%), suggesting that stem/progenitor cells contributes greatly to tHO during this period (Figure 2d). Consequently, we analyzed the differentially expressed genes in the Mesenchymal subcluster at 7 and 21 days post-injury compared with the control group and revealed that 82 genes were upregulated in the Mesenchymal subcluster at Day 7, and 53 genes were upregulated at Day 21 (Figure 2e).

**Figure 2.**
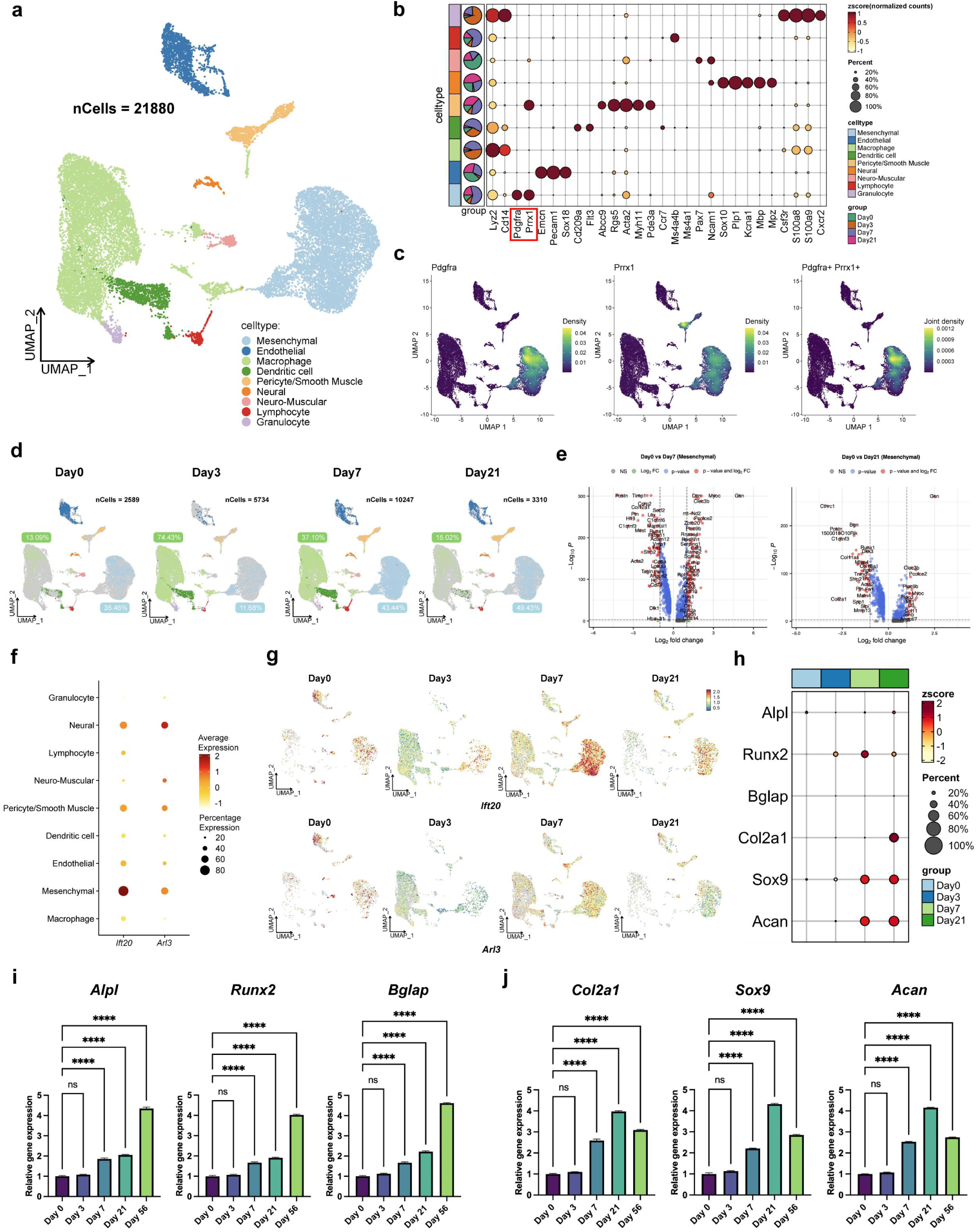
scRNA-seq indicated the role of stem/progenitor cells and ciliary genes in tHO development. (a) UMAP plot of nine clusters identified in 21880 cells in total of tHO injured site. (b) Bubble plot shows the high variable marker genes of each cluster. The red box highlights Pdgfra and Prrx1 as markers for the Mesenchymal cluster. (c) The expression density of Pdgfra+, Prrx1+, and Pdgfra+Prrx1+ cells in each cluster are shown by UMAP plot. The purple-to-yellow gradient signifies a progressive increase in expression levels. (d) UMAP plot of nine clusters with single cells grouped by different time points (Day 0, 3, 7, 21) during tHO demonstrates the trend of changes in cell composition. (e) The volcano plot visualizes the differential expressed genes in the Mesenchymal subcluster at Day 7 (left) or Day 21 (right) compared to the Day 0 group. (f) The dot plot visualizes the different expression of *Ift20* and *Arl3* in nine clusters. (g) UMAP plot shows the expression of *Ift20* (upper) and *Arl3* (lower) at different time points (Day 0, 3, 7, 21) during tHO development. (h) Bubble plot shows the different expression of osteogenic and chondrogenic genes in four groups during tHO development. (i) qPCR showed relative osteogenesis-related genes (*Alpl, Runx2,* and *Bglap*) mRNA expression in different time points (Day 0, 3, 7, 21) during tHO development. (j) qPCR showed relative chondrogenesis-related genes (*Col2a1,Sox9,*and *Acan*) mRNA expression in different time points (Day 0, 3, 7, 21) during tHO development. Data are presented as means ± SD of three independent assays. Statistical analyses were performed by one-way ANOVA analyses with Tukey’s post hoc test for multiple comparisons. ****P < 0.001; ns, P > 0.05.

Since the percentage of ciliated stem/progenitor cells greatly increased during tHO development, we are interested in the expression of ciliary genes at single-cell level. Intraflagellar transport protein 88 homolog (IFT88) functions as a core protein of the IFT-B complex and drives anterograde transport within cilia. Many studies reported the role of IFT88 in skeletal homeostasis^23^ and bone diseases^24–27^. *Ift88* has low expression at single cell level likely due to limited scRNA-seq depth, but still predominantly localizes to the Mesenchymal subpopulation (Figure S2a, b), and shows a subtle upward trend during tHO (Figure S2c). We then shifted our focus to another ciliary gene IFT20 which is also a component of the IFT-B complex and has been reported to regulate ciliogenesis and skeletal development^28^. It can be shown that *Ift20* are highly expressed in the Mesenchymal cluster (Figure 2f). Arf-like Protein 3 (ARL3), a small GTPase, is involved in ciliogenesis and the maintenance of ciliary function^29,30^. Specifically deleting ARL3 in muscle satellite cells impaired muscle hypertrophy, revealing its potential role of regulating primary cilia in stem/progenitor cells. Indeed, *Arl3*mainly expressed in Mesenchymal cluster in cells from tHO injured site (Figure 2f). Moreover, with the development of tHO after injury, the expression of *Ift20* and *Arl3* in Mesenchymal cluster rapidly increased in Day 7 and Day 21 group, while the trends over time were not significant in other clusters (Figure 2g). qPCR analysis further demonstrated that the mRNA expression levels of ciliary genes*Ift88,Ift20*, and *Arl3*had an increasing trend during tHO development and peaked on Day 21 before slightly decreased on Day 56 (Figure S2d). We then explore the expression of osteogenic (*Alpl*, *Runx2*, and *Bglap*) and chondrogenic (*Col1a1,Sox9*, and *Acan*) genes during tHO development. scRNA-seq data showed these genes, especially chondrogenic genes were mainly expressed in Mesenchymal cluster (Figure S2e) and showed increasing trends during tHO progression with a higher expression in Day 7 and Day 21 after injury (Figure 1h). qPCR analysis also showed the expression levels of osteogenic genes gradually increased and reach the peaks at Day 56 (bone formation period) after injury (Figure 1i), while chondrogenic genes peaked at Day 21 (cartilage formation period) and slightly reduced in Day 56 during tHO progression (Figure 1j). These results indicated primary cilia and ciliary genes may mainly functioned in stem/progenitor cells osteochondrogenesis during the progression of tHO.

### 2. Inhibition of primary cilia alleviates TSPCs osteochondrogenesis and tHO

Since the above results suggested that primary cilia on stem/progenitor cells may be involved in tHO pathogenesis, and considering that abnormal differentiation of tendon stem/progenitor cells (TSPCs) has been reported to potentially contribute to the pathogenesis of tHO^31,32^, we isolated TSPCs from the tendon tissues of C57BL/6 mice. Microscopic examination showed that these TSPCs exhibited a fibroblast-like spindle-shaped morphology (Figure 3a). Flow cytometry analysis demonstrated that 88.1% of the TSPCs were double-positive for Pdgfrα and Prrx1 (consistent with the scRNA-seq results), thereby confirming the stem/progenitor cell characteristics of the isolated TSPCs (Figure 3a). Immunofluorescence staining for ARL13B and Acetyl-α-Tubulin (AcTu) was used to observe changes of primary cilia in TSPCs following knockdown of ciliary genes IFT88 and ARL3 using letivirus-shRNA (*shIft88*and *shArl3*). The results showed a significant reduction in cilia counts and length (Figure 3b). Quantitative analysis revealed that the proportion of ciliated TSPCs was approximately 53.54% in the control group but decreased to 19.36% in the *shIft88*group and 34.70% in the *shArl3*group, indicating a marked decrease in cilia number in TSPCs following ciliary gene knockdown (Figure 3c). Additionally, the average cilia length was 3.37 µm in the control group but reduced to 2.06 µm in the *shIft88*group and 2.14 µm in the *shArl3*group, demonstrating a significant shortening of cilia length in TSPCs after ciliary gene knockdown (Figure 3d).

**Figure 3.**
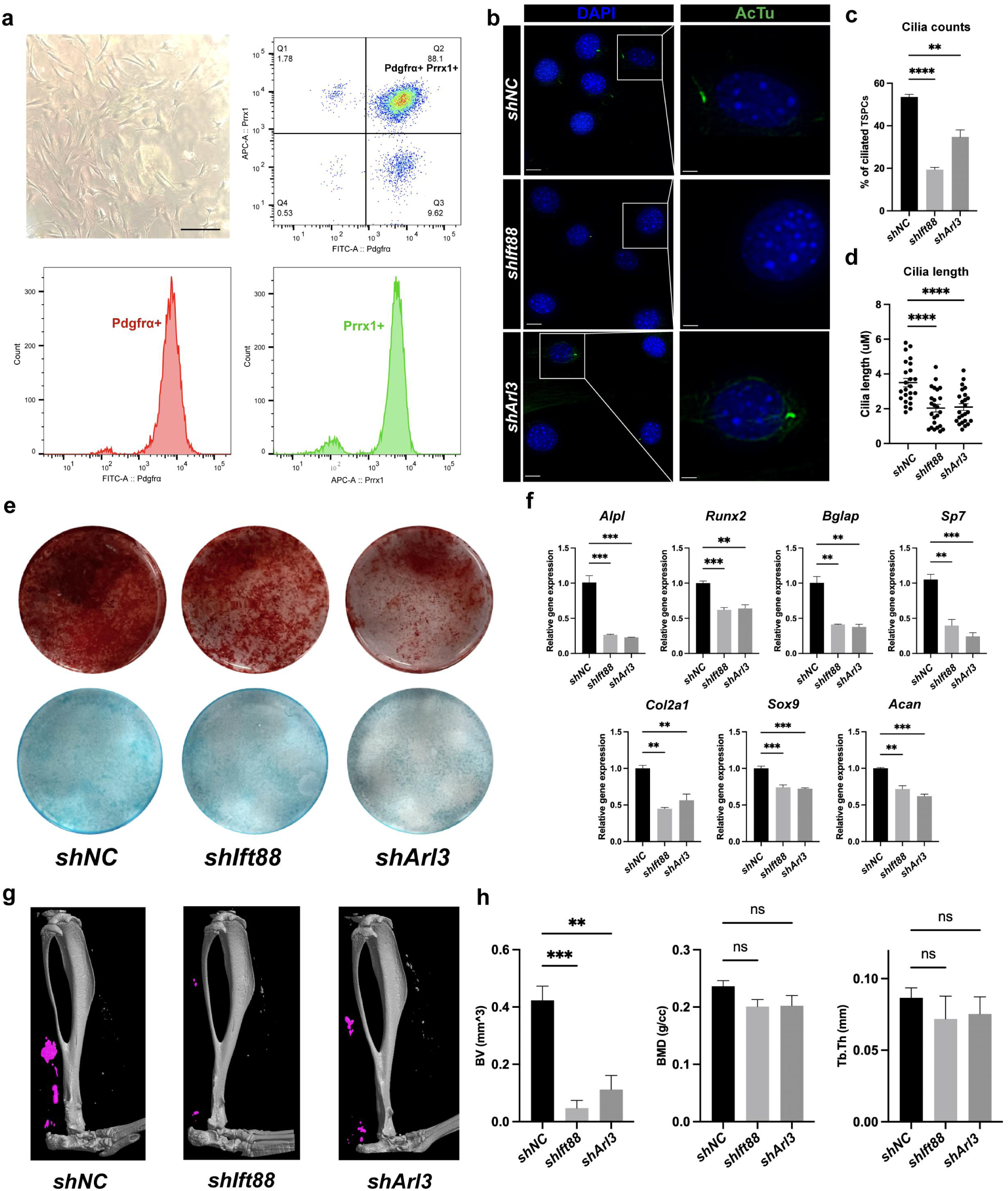
Primary cilia dysfunction impairs TSPCs osteochondrogenesis and ectopic bone formation. (a) Fibroblast-like TSPCs extracted from mice for subsequent experiments (scale bar 100 µm) and flow cytometry analysis of the expression of Pdgfra and Prrx1 (stem/progenitor cell surface markers) on TSPCs. (b) Immunofluorescence images of AcTu (green) mark the cilia in *shNC, shIFT88,* and *shARL3* TSPCs. DAPI staining indicates nuclei (blue), scale bar 30 µm (left) and 10 µm (right). (c) Cilia counts (qualified by the percentage of ciliated TSPCs) in TSPCs with or without *shIFT88* or *shARL3* treatment. (d) Cilia length in TSPCs with or without *shIFT88* or *shARL3* treatment (n= 25/group). (e) Representative images of alizarin red staining (upper) and alcian blue staining (lower) in TSPCs with or without *shIFT88* or *shARL3* treatment during in vitro osteogenic or chondrogenic differentiation. (f) qPCR showed relative osteogenesis-related genes (*Alpl, Runx2, Bglap, and Sp7*) and chondrogenesis-related genes (*Col2a1, Sox9, and Acan*) mRNA expression in TSPCs with or without *shIFT88* or *shARL3* treatment after during in vitro osteogenic or chondrogenic differentiation. (g) Micro-CT images show tHO of lower limbs among *shNC, shIFT88,* and *shARL3* groups (n= 4/group). (h) Bone volume (BV, mm^3), bone mineral density (BMD, g/cc), and Tb.Th (trabecular thickness, mm) are quantified. Independent experiments (n= 4/group) were performed for measurements. Data are presented as means ± SD of three independent assays. Statistical analyses were performed by one-way ANOVA analyses with Tukey’s post hoc test for multiple comparisons. *P < 0.05; **P < 0.01; ***P < 0.005; ****P < 0.001; ns, P > 0.05.

Previous studies have shown that the osteogenic and chondrogenic differentiation capabilities of TSPCs are abnormally activated during the process of tHO, leading to the formation of ectopic bone in tendon tissues^32^. Therefore, investigating whether cilia can regulate the TSPCs osteochondrogenesis is important for exploring the regulatory mechanisms of cilia in tHO. We first induced osteogenic differentiation in TSPCs and performed alizarin red staining. Compared with the control group, the alizarin red staining was lighter in the *shIft88*and *shArl3*groups, indicating less calcium deposition (Figure 3e, S3a). This suggests that knockdown of ciliary genes attenuates the osteogenic differentiation capacity of TSPCs. qPCR analysis revealed that the expression of osteogenesis-related genes *Alpl*, *Bglap*, *Run2*and *Sp7*was significantly reduced in the *shIft88*and *shArl3*groups compared with the control group (Figure 3f). Similarly, we induced chondrogenic differentiation in TSPCs with or without ciliary gene knockdown and performed alcian blue staining. Compared with the control group, the alcian blue staining was lighter in the *shIft88*and *shArl3*groups (Figure 3e, S3a). Meanwhile, qPCR analysis revealed that the expression of chondrogenesis-related genes *Col2a1,Sox9*and *Acan*was significantly reduced (Figure 3f). These findings identified a positive correlation between cilia function and the osteochondrogenic differentiation capacity of TSPCs.

To locally inhibit ciliary function in vivo, we administered adeno-associated virus (AAV) vectors carrying *shIft88* and *shArl3* via intramuscular injection into the lower limbs of mice with tHO. The results showed that knockdown of*Ift88*and *Arl3*significantly decreased the bone volume (BV) of ectopic bone formation, whereas bone mineral density (BMD) and trabecular thickness (Tb.Th) was not significantly altered (Figure 3g, h). These results indicate that primary cilia may positively modulate the osteogenic and chondrogenic differentiation of TSPCs, leading to abnormal bone formation in local soft tissues and ultimately resulting in tHO, which could be rescued by inhibition of ciliary function without influencing normal bone quality.

### 3. Ciliary Hh and PI3K-Akt signaling are involved in tHO pathogenesis

To further investigate the specific molecular mechanisms and downstream targets through which primary cilia regulate tHO development, we conducted enrichment analysis on our previously mentioned sequencing data. Gene Ontology (GO) enrichment analysis showed that differentially expressed genes at the site of tHO 21 days post-injury were significantly enriched in functions related to the extracellular matrix, ossification, and cartilage development (Figure 4a). Consistent with these findings, differentially expressed genes in TSPCs following knockdown of ciliary genes were also enriched in skeletal system development, ossification, and bone mineralization (Figure 4b). These results suggest a strong correlation between the differentially expressed genes and tHO, indicating that targeting these genes may represent a potential strategy for treating tHO. We further conducted Gene Set Enrichment Analysis (GSEA) on the differentially expressed genes in TSPCs following knockdown of ciliary gene *Ift88*, and the results showed significant enrichment and negative correlation (NES = -2.124) in the Hh signaling pathway (Figure 4c). Given that the activity of the Hedgehog (Hh) signaling pathway is dependent on the primary cilia for regulating embryonic development and tissue differentiation, and considering that aberrant upregulation of Hh has been implicated in the pathogenesis of tHO following injury^33^, the Hh signaling activity within TSPCs cilia is likely closely related to tHO. Additionally, KEGG enrichment analysis indicated that differentially expressed genes in the Mesenchymal cluster of tHO tissues, as well as those in TSPCs following knockdown of ciliary gene*Ift88*, were significantly enriched in the PI3K-Akt signaling pathway (Figure 4d, e).

**Figure 4.**
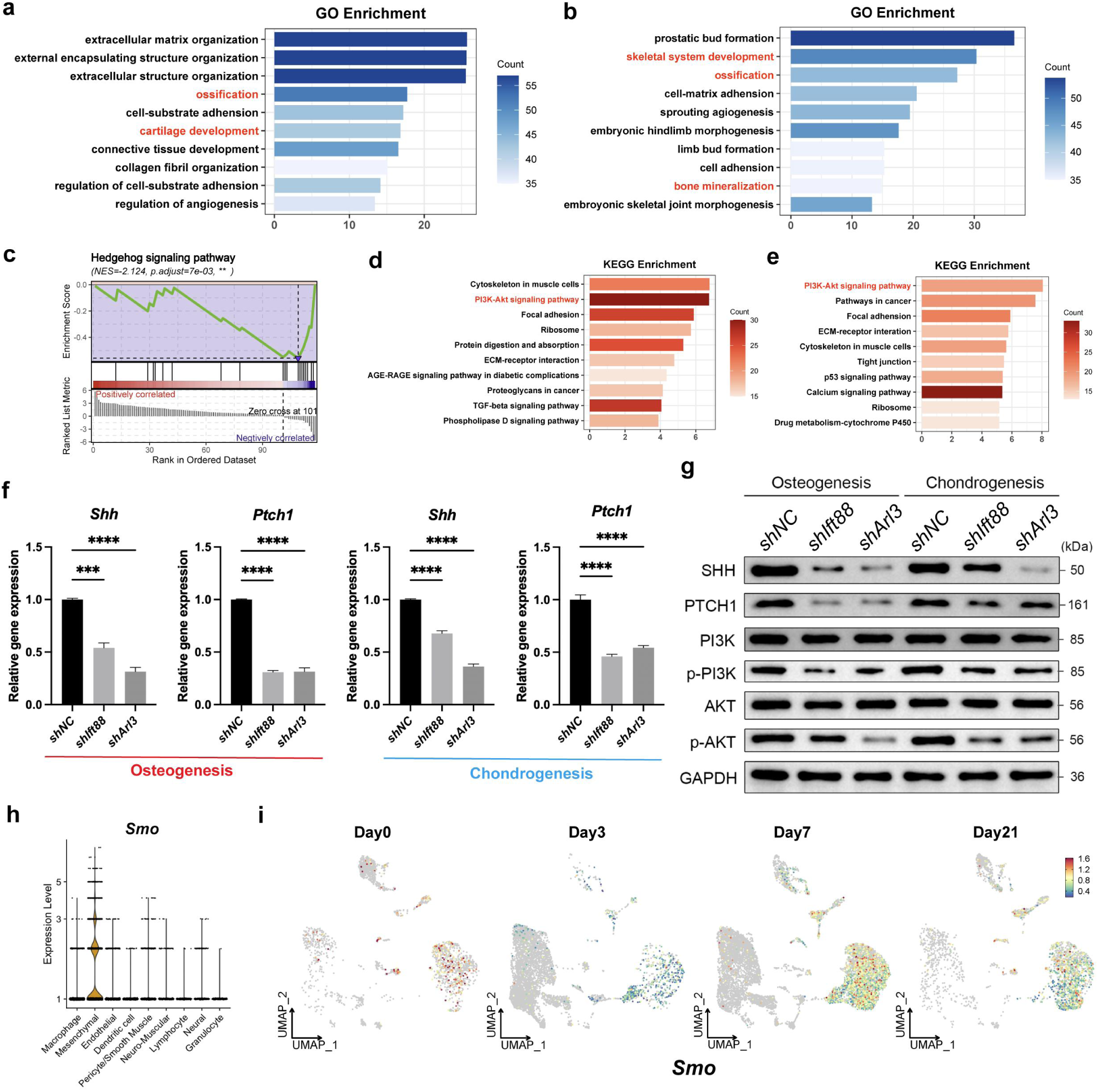
TSPCs ciliary Hh and PI3K-Akt signaling are involved in tHO progression. (a) GO enrichment analysis of differential expressed genes in tHO mice injured site shows top 10 enriched biological processes that are important for tHO. (b) GO enrichment analysis of differential expressed genes in TSPCs treated with *shIft88* shows top 10 enriched biological processes. (c) GSEA enrichment of TSPCs treated with *shIft88* compared with *shNC*. The enriched pathway with the absolute value of normalized enrichment score (NES) greater than 1 and adjusted P-value less than 0.05 was considered significant. (d) KEGG enrichment analysis of differential expressed genes in Mesenchymal subcluster of tHO injured site identified by scRNA-seq shows top 10 enriched pathways. (e) KEGG enrichment analysis of differential expressed genes in TSPCs treated with *shIft88* showed top enriched pathways. (f) qPCR showed relative Hh signaling genes (Shh, Ptch1) mRNA expression in TSPCs with or without *shIft88* or *shARL3* treatment during in vitro osteogenic or chondrogenic differentiation. (g) Western blot showed Hh signaling (SHH, PTCH1) and PI3K-Akt signaling (PI3K, p-PI3K, AKT, p-AKT) protein expression levels in TSPCs with or without *shIFT88* or *shARL3* treatment during in vitro osteogenic or chondrogenic differentiation. (h) The expression level of *Smo* in each cluster are shown by violin plot. (i) UMAP plot shows the expression of *Smo* at different time points (Day0, 3, 7, 21) during tHO development. Data are presented as means ± SD of three independent assays. Statistical analyses were performed by one-way ANOVA analyses with Tukey’s post hoc test for multiple comparisons. ***P < 0.005; ****P < 0.001

To verify the bioinformatics analysis results, we performed qPCR analysis and found that the mRNA expression of Hh signaling molecules *Shh*and *Ptch1*was significantly downregulated in TSPCs undergoing osteogenic and chondrogenic differentiation after *shIft88*or *shArl3*treatment (Figure 4f). Western blot analysis also demonstrated a significant reduction in the protein expression levels of SHH and PTCH1 (Figure 4g). Additionally, during TSPCs osteochondrogenesis, the ratios of phosphorylated PI3K to total PI3K (p-PI3K/PI3K) and phosphorylated AKT to total AKT (p-AKT/AKT) were significantly decreased following ciliary gene knockdown (Figure 4g), indicating reduced phosphorylation of PI3K and AKT and the inactivation of the PI3K-Akt signaling. Smoothened (SMO) is a key seven-transmembrane receptor involved in Hh signaling and is inhibited by PTCH1 when Hh signaling is inactive. Upon binding of the Hh ligand to PTCH1, the inhibitory effect of PTCH on SMO is alleviated, thereby enabling the activation of SMO. Interestingly, scRNA-seq analysis showed that *Smo* mainly expressed (Figure 4h) and increased over time in Mesenchymal cluster (Figure 4i) during tHO progression, further demonstrating the Hh signaling was activated in tHO. Collectively, these results suggest that the primary cilia of TSPCs may lead to aberrant activation of Hh and PI3K-Akt signaling, thereby promoting TSPCs osteochondrogenesis and exacerbating tHO development.

### 4. Primary cilia modulate MEST expression to promote tHO

To further identify key genes involved in tHO and their potential crosstalk with primary cilia, we obtained bulk-RNA sequencing data and analyzed the differentially expressed genes (Figure S4a) from a public database for mouse tHO tissues (GSE233201). We then integrated these data with RNA sequencing results from TSPCs with ciliary gene *Ift88* knockdown and control groups (Figure S4b) to identify common differentially expressed genes. Genes upregulated in the Mesenchymal cluster at 7 and 21 days during tHO (GSE126060), genes upregulated in injured tissue at 21 days of tHO (GSE233201), and genes downregulated in TSPCs with ciliary gene *Ift88* knockdown were obtained respectively (Table S1). Venn diagram analysis identified three common differentially expressed genes: MEST, PTN, and COL2A1 among these three datasets (Figure 5a). COL2A1 is a key gene for chondrogenic differentiation, while PTN showed insignificant differences after validation (Figure S5a-d). Thus, we focused on the role of the MEST gene in tHO in subsequent studies.

**Figure 5.**
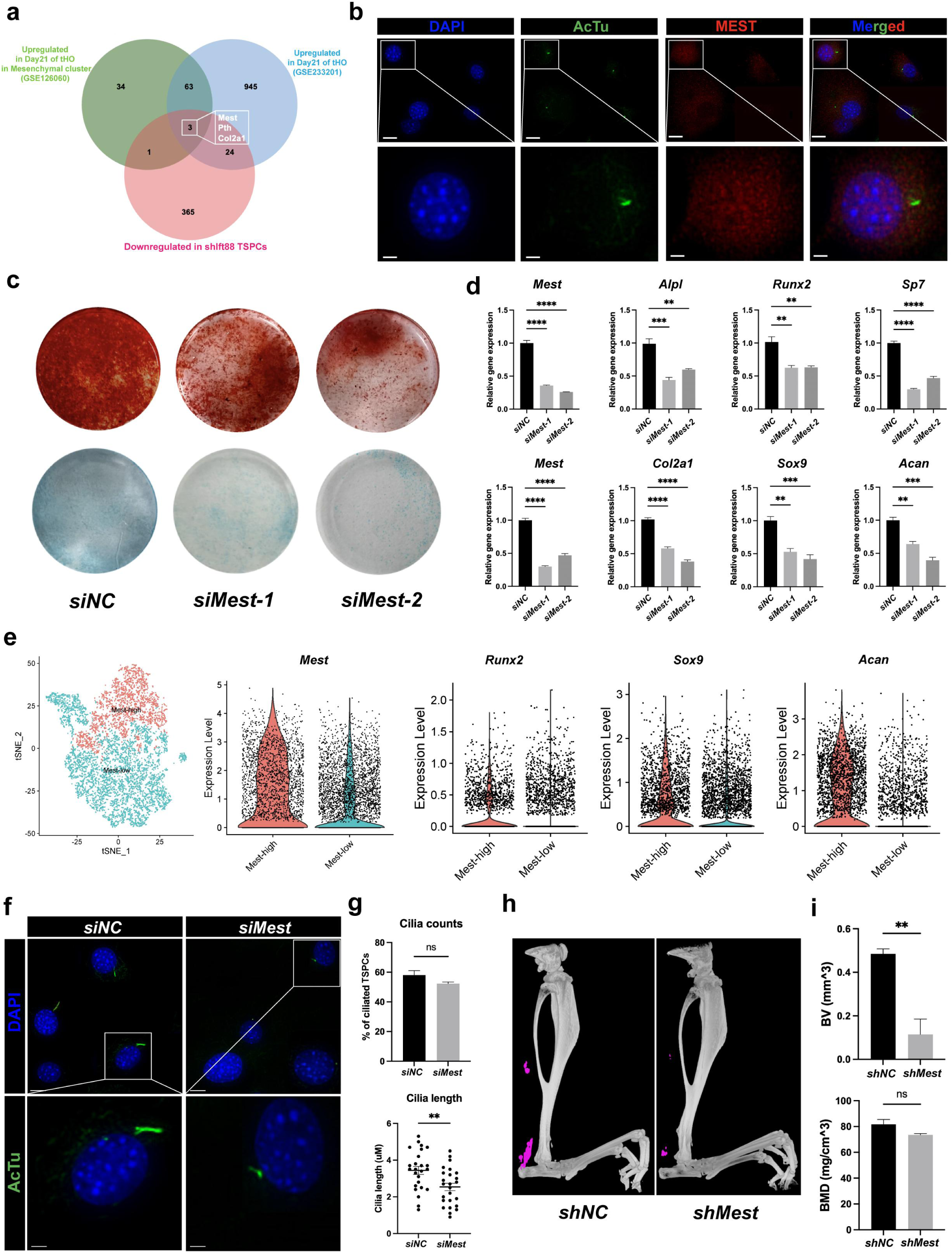
MEST modulates TSPCs cilia length and osteochondrogenesis in tHO. (a) Venn diagram shows the common differential expressed genes among three gene sets and identified MEST, PTN, COL2A1 as the key genes. (b) Immunofluorescence images of AcTu (green) and MEST (red) in TSPCs. DAPI staining indicates nuclei (blue), scale bar 30 µm (upper) and 10 µm (lower). (c) Representative images of alizarin red staining (upper) and alcian blue staining (lower) in TSPCs with or without *siMest* treatment during in vitro osteogenic or chondrogenic differentiation. (d) qPCR shows relative *Mest* and osteogenesis-related (*Alpl, Runx2, Bglap*) or chondrogenesis-related genes (*Col2a1, Sox9, Acan*) mRNA expression in TSPCs with or without *siMest* treatment during in vitro osteogenic or chondrogenic differentiation. (e) tSNE map shows Mest-high and Mest-low subclusters divided from Mesenchymal cluster. The expression level of *Mest, Runx2, Sox9*, and *Acan* in Mest-high and Mest-low subclusters were visualize by violin plots. (f) Immunofluorescence images of AcTu (green) mark the cilia of TSPCs. DAPI staining indicates nuclei (blue), scale bar 30 µm (upper) and 10 µm (lower). (g) Cilia counts (qualified by ciliated TSPCs proportion) and cilia length (n= 25/group) in the *siNC* and *siMest* group. (h) Micro-CT images show tHO of lower limbs in *shNC* and *shMest* groups. (i) Quantification of tHO degree by bone volume (BV, mm^3) and bone mineral density (BMD, mg/cm^3). Independent experiments (n= 4/group) were performed for measurements. Data are presented as means ± SD of three independent assays. Statistical analyses were performed by Student’s t-test for two-group comparison and one-way ANOVA analyses with Tukey’s post hoc test for multiple comparisons. *P < 0.05; **P < 0.01; ***P < 0.005; ****P < 0.001; ns, P > 0.05.

scRNA-seq analysis revealed that *Mest*is predominantly expressed in the Mesenchymal cluster (Figure S4c, d). Both qPCR and Western blot analyses demonstrated a significant reduction in *Mest*RNA and protein expression following knockdown of the ciliary genes*Ift88*or *Arl3*(Figure S4e, f). Moreover, scRNA-seq analysis showed that the expression of *Mest* in the Mesenchymal cluster exhibits an upward trend with the progression of tHO (Figure S4g). Consistently, immunohistochemical analysis of tendon tissues at various time points after establishment of tHO further showed that MEST protein expression is progressively upregulated during the tHO development (Figure S4h), suggesting that MEST may play a significant role in the regulatory mechanisms of tHO. Notably, immunofluorescence staining showed no significant co-localization between MEST and AcTu (Figure 5b), indicating that MEST likely functions outside but not within the cilia in TSPCs to influence cilia function and thereby contribute to the occurrence and progression of tHO.

### 5. Targeting MEST inhibits TSPCs osteochondrogenesis and tHO formation

As MEST is primarily not localized within the cilia of TSPCs, we explored how MEST can regulate tHO in a cilia localization-independent manner. We constructed two small interfering RNAs (siRNAs): *siMest-1*and *siMest-2*, to inhibit *Mest*gene expression in vitro. qPCR analysis revealed a significant decrease in *Mest*expression following *siMest*intervention, confirming that *siMest*can effectively suppress MEST function (Figure 5d). Alizarin red staining showed that the *siMest*group exhibited lighter staining and fewer calcium nodules compared with the control group during osteogenic differentiation induction (Figure 5c, S3b). Additionally, the expression of osteogenesis-related genes *Alpl,Runx2*and *Bglap* was significantly reduced (Figure 5d). Similarly, alcian blue staining during chondrogenic differentiation induction showed that, compared with the control group, the *siMest*group exhibited lighter staining and reduced chondrogenic capacity (Figure 5c, S3b). The expression of chondrogenesis-related genes *Col2a1*, *Sox9*and *Acan*was also decreased (Figure 5d). Moreover, we divided the Mesenchymal cluster to Mest-high and Mest-low subclusters based on the expression level of Mest in burn/tenotomy tHO scRNA-seq dataset (Figure 5e). Though *Alpl, Sp7,* and *Col2a1* were not significantly expressed between two subclusters probably because of their extreme low expression in single-cell level (Figure S5e), the expression of *Runx2,Sox9,*and *Acan*were greatly higher in Mest-high subcluster (Figure 5e). These findings indicated that alleviating MEST expression diminish the osteochondrogenic differentiation capacity of TSPCs.

We further employed immunofluorescence staining with AcTu to label the cilia of TSPCs in the control and *siMest*groups (Figure 5f). Quantitative analyses (Figure 4g) showed no significant difference in cilia number (58.26% vs 53.11%), while cilia length was significantly shorter in the *siMest*group compared with the control group (2.46 µm vs 3.48 µm). In addition, we performed in vivo experiments by locally injecting a AAV vector carrying *shMest*into mice with tHO to silencing *Mest*expression in local injured tissues. Micro-CT imaging revealed that ectopic bone volume in the *shMest*group was significantly lower than that in the control group, while bone mineral density remained largely unchanged (Figure 5h, i). Collectively, these results suggest that MEST can regulate ciliary function and TSPCs osteochondrogenesis in a ciliary localization-independent manner, while targeting MEST may serve as a potential method to restore normal TSPCs osteochondrogenesis and treat tHO.

### 6. Activation of MEST transcription by GLI2 mediates Hh signaling in tHO

GLI2, a key transcriptional activator of the Hh signaling pathway, can translocate to the nucleus upon Hh signal activation and induces the expression of downstream genes involved in cell proliferation and differentiation. The sequence logo of GLI2 transcription factor was shown and the JASPAR database predicts that the GLI2 can bind to the MEST gene promoter at positions 904-916 (gtgaccacccagc) and 1698-1710 (tggaccacgcact), both with relative score > 0.8 (Figure 6a). Dual-luciferase reporter assays using pGL4.10 as vector demonstrated that GLI2 can bind to the MEST promoter at both predicted sites, with overexpression of GLI2 significantly increasing luciferase activity, thus indicating that GLI2 positively regulates MEST transcription (Figure 6b). To further elucidate the regulatory role of the GLI2 transcription factor in tHO, we performed immunohistochemistry on injured tendon tissues obtained from mice with tHO and found that GLI2 expression progressively increased during tHO development (Figure 6c). Meanwhile, qPCR and Western blot analyses revealed that, compared with the control group, the *Gli2*mRNA and protein expression levels of GLI2 were significantly downregulated with *shIft88* or *shArl3*treatment during TSPCs osteochondrogenesis (Figure 6d, e). ChIP-qPCR assay further demonstrated that GLI2 transcriptional factor could significantly bind to site 1/2 on MEST promoter region (Figure 6f). To further investigate the role of GLI2 in regulating MEST transcription, overexpression (GLI2-OE) and knockdown (*siGli2*) of Gli2 were performed in TSPCs, affecting the Mest mRNA expression during both osteogenic and chondrogenic differentiation, which further demonstrated that GLI2 activity was positively associated with MEST gene transcription in TSPCs osteochondrogenesis (Figure 6g). Moreover, local injection of AAV-*shMest*into tHO mice injured site decreased protein expression levels of SHH, GLI2, PTCH1, and SMO, as shown by immunohistochemistry (Figure 6h). qPCR analysis also showed reduced mRNA expression of Hh signaling molecules (*Shh, Ptch1, Gli2*, and *Smo)* in *shMest* treated tHO mice (Figure 6i). Thus, the in vivo experiments indicated that targeting Mest may alleviate the occurrence and progression of tHO by reducing the activation level of the Hh/Gli2 signaling. Collectively, these results suggested GLI2 as a positive regulator of MEST transcription, while MEST in turn modulates GLI2 and Hh signaling activity during tHO development.

**Figure 6.**
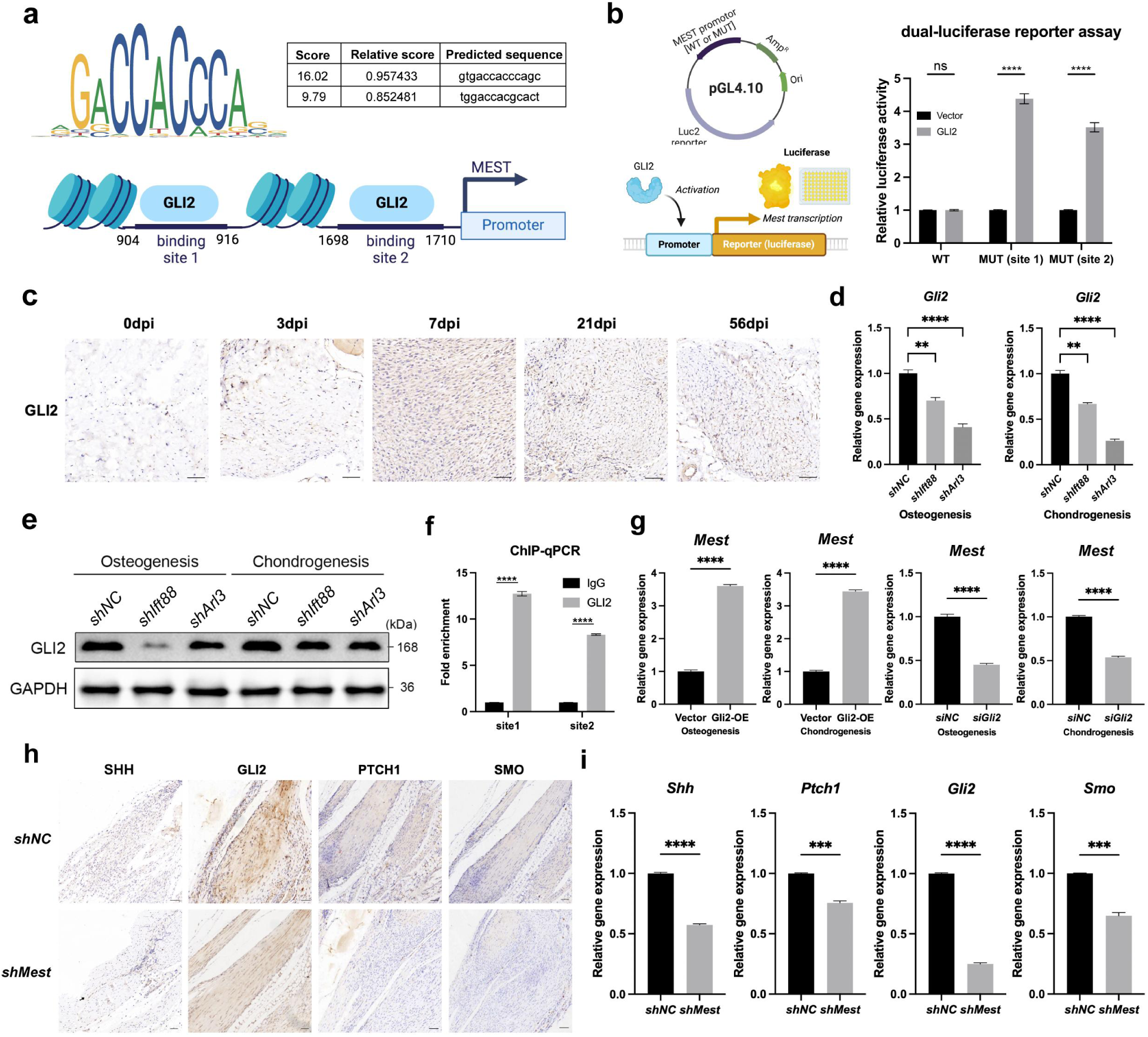
GLI2 binds to the MEST promoter to activate its transcription via Hh signaling. (a) The sequence logo of GLI2 transcription factor and two potential binding sites of the GLI2 transcription factor on the MEST promoter region predicted by the JASPAR database. (b) The luciferase reporter assay using pGL4.10 as vector showed that GLI2 can bind to the MEST promoter at both site 1 and 2. (c) Immunohistochemistry images demonstrated GLI2 in tHO tendon tissues at different time post injury (scale bar 100 µm). (d) qPCR shows relative *Gli2* mRNA expression in TSPCs with or without *shIft88* or *shARL3* treatment during in vitro osteogenic or chondrogenic differentiation. (e) Western blot shows GLI2 protein expression levels in TSPCs with or without *shIFT88* or *shARL3* treatment during in vitro osteogenic or chondrogenic differentiation. (f) ChIP-qPCR shows GLI2 can binds to both the site 1 and site 2 of the MEST promoter region. (g) qPCR showed relative Mest mRNA expression in TSPCs with or without Gli2-OE or *siGli2*treatment during in vitro osteogenic or chondrogenic differentiation. (h) Immunohistochemistry images demonstrated SHH, GLI2, PTCH1, and SMO expression in tHO tendon tissues with *shNC* or *shMest* treatment after injury (scale bar 150 µm). (i) qPCR shows relative *Shh, Ptch1, Gli2,* and *Smo* mRNA expression in tHO tendon tissues with *shNC* or *shMest* treatment after injury. Data are presented as means ± SD of three independent assays. Statistical analyses were performed by Student’s t-test for two-group comparison and one-way ANOVA analyses with Tukey’s post hoc test for multiple comparisons. **P < 0.01; ***P < 0.005; ****P < 0.001; ns, P > 0.05.

### 7. MEST regulates TSPCs osteochondrogenesis via Hh and PI3K-Akt signaling

Previous results have confirmed that: 1) MEST can influence TSPCs osteochondrogenesis and regulate tHO development in ciliary localization-independent manner; 2) GLI2 promote MEST transcription during TSPCs osteochondrogenesis via Hh signaling; 3) Cilia-mediated Hh and PI3K-Akt signaling play a crucial role in tHO pathogenesis. Thus, we became intrigued by whether MEST regulates TSPCs osteochondrogenesis through Hh and PI3K-Akt signaling. We first transfected *siMest*in TSPCs to knock down Mest gene expression in vitro. Compared with *siNC*treated TSPCs, the expression levels of key Hh signaling molecules were downregulated at both the mRNA (Figure 7a) and protein (Figure 7b) levels in *siMest*treated TSPCs during osteogenic and chondrogenic differentiation. In addition, the phosphorylation levels of PI3K and AKT were also decreased significantly (Figure 7c). These findings suggest that silencing Mest expression can attenuate TSPCs osteochondrogenesis via Hh and PI3K-Akt signaling in tHO.

**Figure 7.**
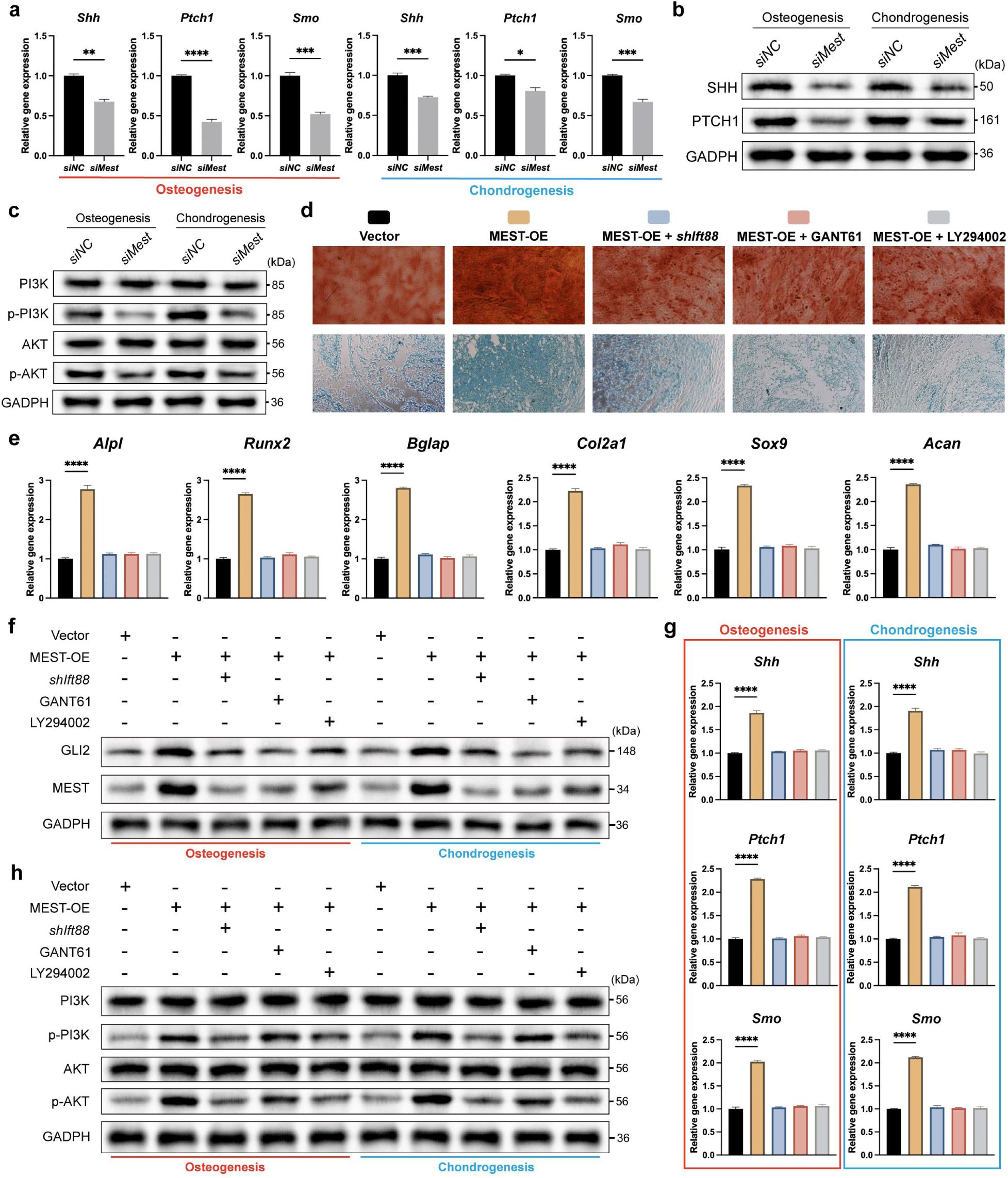
MEST enhances TSPCs osteochondrogenesis through Hh and PI3K-Akt signaling. (a) qPCR showed relative Hh signaling genes (*Shh, Ptch1, and Smo*) mRNA expression in TSPCs with or without *siMest* treatment during in vitro osteogenic or chondrogenic differentiation. (b) Western blot shows Hh signaling (SHH, PTCH1) protein expression levels in TSPCs with or without *siMest* treatment during in vitro osteogenic or chondrogenic differentiation. (c) Western blot showed PI3K-Akt signaling (PI3K, p-PI3K, AKT, p-AKT) protein expression levels in TSPCs with or without *siMest* treatment during in vitro osteogenic or chondrogenic differentiation. (d) Representative images of alizarin red staining (upper) and alcian blue staining (lower) in TSPCs during in vitro osteogenic (upper) and chondrogenic (lower) differentiation in Vector, MEST-OE, MEST-OE + *shIft88*, MEST-OE + GANT61, and MEST-OE + LY294002 treated groups. (e) qPCR showed relative osteogenesis-related genes (*Alpl, Runx2*, *Bglap*) and chondrogenesis-related genes (*Sox9,Col2a1,Acan*) mRNA expression after in vitro TSPCs osteogenic or chondrogenic differentiation in five groups. (f) Western blot showed GLI2 and MEST protein expression levels in TSPCs under different treatment during in vitro osteogenic or chondrogenic differentiation. (g) qPCR showed relative Hh signaling genes (*Shh, Ptch1, and Smo*) mRNA expression in TSPCs under different treatment during in vitro osteogenic or chondrogenic differentiation. (h) Western blot showed PI3K-Akt signaling (PI3K, p-PI3K, AKT, p-AKT) protein expression levels in TSPCs under different treatment during in vitro osteogenic or chondrogenic differentiation. Data are presented as means ± SD of three independent assays. Statistical analyses were performed by Student’s t-test for two-group comparison and one-way ANOVA analyses with Tukey’s post hoc test for multiple comparisons. *P < 0.05; **P < 0.01; ***P < 0.005; ****P < 0.001.

To further demonstrate whether: 1) The regulatory effects of primary cilia on TSPCs osteochondrogenesis and tHO development is dependent on MEST activity; 2) Hh or PI3K-Akt signaling is essential for MEST-induced hyperactivation of TSPCs osteochondrogenesis, we performed rescue experiments in MEST overexpression (MEST-OE) TSPCs by targeting cilia or Hh/PI3K-Akt signaling. Both osteogenic and chondrogenic differentiation were upregulated in MEST-OE group, indicated by higher positive area of alizarin red and alcian blue staining (Figure 7d, S3c). However, the effects can be significantly rescued by inhibiting cilia function under *shIft88*treatment in MEST-OE TSPCs (Figure 7d, S3c), which reveals that primary cilia promote TSPCs osteochondrogenesis by activating MEST. Targeting Hh and PI3K-Akt signaling by GANT61 (GLI2 inhibitor) and LY294002 (PI3K inhibitor) respectively in MEST-OE TSPCs also restored the normal osteochondrogenesis (Figure 7d, S3c), suggesting that Hh and PI3K-Akt signaling serve as critical mediators through which MEST enhances TSPCs differentiation. qPCR analysis showed consistent changes of osteogenesis (*Alpl,Runx2,*and *Bglap*) and chondrogenesis (*Col2a1,Sox9,*and *Acan*) related genes expression with the staining results under different treatment (Figure 7e). We also found that overexpression of MEST (MEST-OE) lead to upregulated GLI2 (Figure S3d, 7f) and other Hh signaling molecules (*Shh,Ptch1*, and *Smo*) expression (Figure 7g), as well as the phosphorylation levels of PI3K and AKT (Figure 7h), which is opposite to the MEST knockdown effects during TSPCs osteochondrogenesis. These findings add to evidence of the crucial role of Hh and PI3K-Akt signaling in MEST-regulated TSPCs osteochondrogenesis and tHO development.

Since both Hh and PI3K-Akt signaling are involved in tHO pathogenesis, exploring the relationship and crosstalk between them is of great significance. qPCR (Figure S3d) and western blot (Figure 7f) showed that GANT61 (the specific inhibitor of GLI2 expression and GLI2-mediated transcription) can reduced both GLI2 and MEST expression in MEST-OE TSPCs during in vitro osteochondrogenesis. However, GANT61 cannot rescue the phosphorylation levels of PI3K and AKT during MEST-OE TSPCs osteochondrogenesis (Figure 7h), while LY294002 (PI3K inhibitor) can significantly attenuate the expression of GLI2/MEST (Figure S3d, 7f) and Hh signaling molecules (Figure 7g), indicating that GLI2 and GLI2-regulated MEST transcription is the downstream of PI3K/Akt signaling and the activation of GLI2-MEST axis involved in TSPCs osteochondrogenesis can be inhibited by targeting PI3K/Akt signaling.

### 8. MEST reduced ciliary sphingomyelin levels in TSPCs

Recent studies have shown that ciliary lipids play a crucial role in maintaining ciliary structure stability and regulating ciliary mechanosensation and signal transduction. Aberrant distribution and dynamic changes in ciliary lipids have been implicated in various diseases, including polycystic kidney disease^17^ and obesity^34^. Moreover, deficiencies in key cholesterol-metabolizing enzymes have been shown to cause ciliary defects that impact bone development^19^. These findings have prompted our interest in investigating the changes in lipids within TSPCs cilia during tHO. We first isolated cilia from TSPCs in the control and *siMest*groups using ultracentrifugation to prepare ciliary suspensions as previously reported^35^ (for more details see Methods). Immunofluorescence showed almost no cilia was left after isolation in non-ciliated TSPCs (Figure 8a). Immunoblot further revealed ciliary enrichment of AcTu and absence of cytoplasmic components (β-actin) in TSPCs cilia (Figure 8b), confirming that the cilia were effectively removed from TSPCs without detectable components of cytoplasm. Then we conducted untargeted lipidomics using liquid chromatography-tandem mass spectrometry (LC-MS/MS) on the collected ciliary suspensions. OPLS-DA analysis showed obvious differences in the lipid profiles of TSPCs cilia between the *siNC* and *siMest*groups (Figure 8c). We performed differential analysis of lipids based on the LIPID MAPS® classification system and the differential expressed primary (Figure 8d, e) and secondary (Figure S6a) lipids were classified. Totally, we screened 159 upregulated and 157 downregulated lipids in the *siMest*group with the criteria of |log2FC| > 1 and P value < 0.05 (Table S2, Figure 8f, S6b). Compared with the *siNC* group, the abundance of sphingomyelin (SM) within the cilia of TSPCs in the *siMest* group was significantly increased, while the content of triglycerides (TG) was significantly decreased (Figure 8g). Heatmaps revealed consistent differential changes in almost all subclasses of SM and TG, including 22 types of SM and 53 types of TG (Figure S6c). Other sphingolipids except for SM also differentially expressed (mostly upregulated) in *siMest* group (Figure S6d). We also conducted KEGG enrichment analysis to find the key lipid signaling involved in *siMest*TSPCs. Consistent with our prior findings, the differentially expressed metabolites were enriched in pathways including glycerophospholipid metabolism, sphingolipid signaling, and sphingolipid metabolism (Figure 8h, S6e). Since SM is the most abundant sphingolipid and exhibits the most significant difference before and after *siMest*knockdown, we then concentrate on the role of ciliary SM metabolism in tHO pathogenesis.

**Figure 8.**
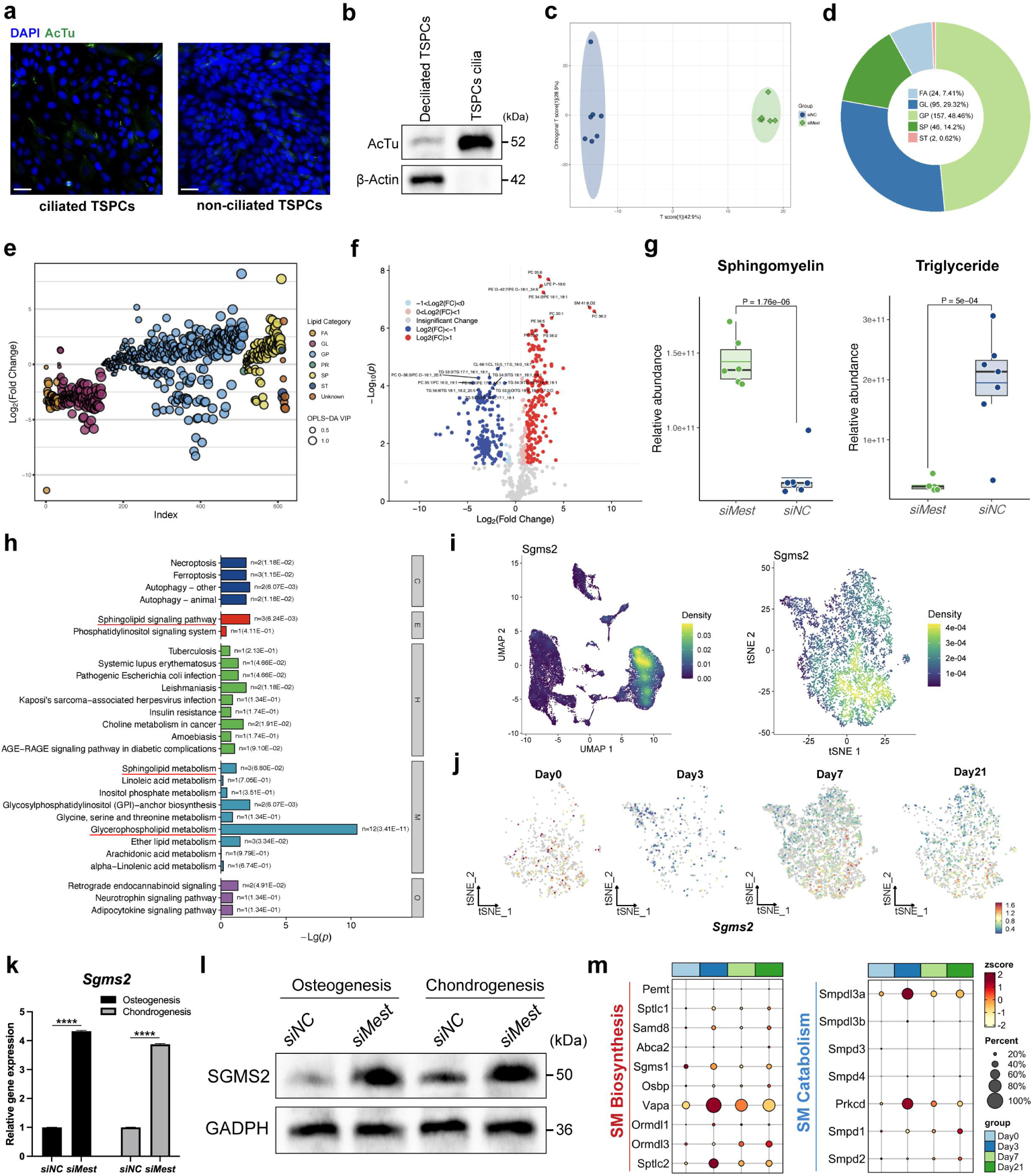
MEST regulates ciliary sphingomyelin metabolism in TSPCs. (a) Immunofluorescence images of AcTu (green) marked cilia in ciliated and non-ciliated mouse primary TSPCs after cilia isolation. DAPI staining indicates nuclei (blue). scale bar 100 µm. (b) Western blot analysis of extracted TSPCs cilia lysates and non-ciliated TSPCs. (c) OPLS-DA plot shows the differences between *siNC* (blue, n= 7) and *siMest* (green, n= 6) groups. (d) Circular diagram of the classification statistics of differential lipids. (e) Bubble chart of lipid classification. The horizontal axis represents the quantity range of lipids, and the vertical axis represents the logarithmic transformation of fold change. Each circle represents a lipid substance. The larger the circle, the higher the variable importance. Different colors represent the primary classification, and lipids of the same classification are arranged together. (f) The volcano plot visualizes the differential expressed lipids in *siMest* group compared to the *siNC* group. Differential expressed lipids are visualized as red (upregulated) or blue (downregulated) bubbles, identified by expression fold-change and p-value. (g) Relative abundance of sphingomyelin and triglyceride are significantly different in *siMest* and *siNC* group. (h) KEGG enrichment analysis of differential expressed lipids between *siMest* and *siNC* group (C: cellular processes; E: environmental information processing; H: human diseases; M: metabolism; O: organismal systems). (i) The expression density of *Sgms2* in each cluster (left) and in Mesenchymal subcluster (right) are shown by UMAP plot. (j) UMAP plot shows the expression of *Sgms2* in Mesenchymal subcluster at different time points (Day0, 3, 7, 21) during tHO development. (k) qPCR shows relative *Sgms2*mRNA expression in TSPCs with or without *siMest*treatment during in vitro osteogenic or chondrogenic differentiation. (l) Western blot shows SGMS2 protein expression levels in TSPCs with or without *siMest*treatment during in vitro osteogenic or chondrogenic differentiation. (m) Bubble plot shows the different expression of SM biosynthesis (red) or catabolism (blue) related genes at different time points (Day 0, 3, 7, 21) during tHO development. Data are presented as means ± SD of three independent assays. Statistical analyses were performed by Student’s t-test for two-group comparison. ****P < 0.001.

SGMS2 (sphingomyelin synthase 2) gene encodes sphingomyelin synthase 2, which is mainly involved in the synthesis of SM on the cell membrane. It has been reported that SGMS2-mediated SM metabolism was involved in skeletal homeostasis regulation and its mutation could lead to a series of bone diseases like osteoporosis and skeletal dysplasia^36^. We found that *Sgms2* was mainly expressed in Mesenchymal cluster but rarely expressed in Mest-high subcluster, while the expression level of *Sgms2*in Mest-high subcluster gradually reduced over time after tHO at single-cell level (Figure 8i, j), indicating that MEST may reduce ciliary SM synthesis in tHO by inhibiting Sgms2 expression. Moreover, qPCR (Figure 8k) and Western blot (Figure 8l) demonstrated that the mRNA and protein expression levels of SGMS2 significantly increased after *siMest*treatment during TSPCs osteogenesis and chondrogenesis. scRNA-seq analysis demonstrated a subtle decrease in SM biosynthetic gene expression during tHO development, while SM catabolic genes showed no significant expression changes (Figure 8m). These results indicated that MEST may reduce the synthesis and abundance of SM within the cilia of TSPCs in tHO progression, and that silencing *Mest*expression can promote SM synthesis by upregulating SGMS2 activity and reverse the reduction of ciliary SM levels in TSPCs.

### 9. Targeting ciliary sphingomyelin attenuates tHO development

Since previous findings have demonstrated that MEST can activate TSPCs osteochondrogenesis in ciliary localization-independent manner, thereby promoting the occurrence and development of tHO, it is essential to explore whether ciliary lipids of TSPCs also contribute to tHO pathogenesis. Research has shown that sphingomyelin and cholesterol interact on the cellular membrane to regulate intracellular signaling processes, and their dynamic balance is crucial for maintaining normal cellular functions^37^. Thus, exploring how MEST-mediated sphingomyelin changes regulate TSPCs function is of great significance in tHO pathogenesis. We induced osteogenic and chondrogenic differentiation in TSPCs and performed alizarin red (ARS) and alcian blue (AB) staining with different treatments respectively (Figure 9a, b). Compared with the control group, treatment with exogenous sphingomyelin significantly reduced the staining intensity and area, indicating that sphingomyelin can suppress TSPCs osteochondrogenesis. Conversely, treatment of TSPCs with myriocin (a sphingomyelin synthesis inhibitor) significantly enhanced TSPCs osteochondrogenesis. However, when sphingomyelin and myriocin were applied together, the ARS or AB staining intensity and area were not significantly different from the control group. Furthermore, downregulation of *Mest*gene expression in TSPCs using *siMest*rescued the excessive upregulation of osteochondrogenesis induced by myriocin treatment in TSPCs. qPCR analysis of the expression of osteogenic (*Alpl, Runx2,*and *Bglap*) and chondrogenic (*Col2a1,Sox9,*and *Acan*) differentiation-related genes in TSPCs revealed trends consistent with the staining results (Figure 9c), suggesting that inhibiting *Mest*gene expression may increase ciliary sphingomyelin level in TSPCs, thereby attenuating their osteogenic and chondrogenic differentiation capacities and alleviating tHO severity.

**Figure 9.**
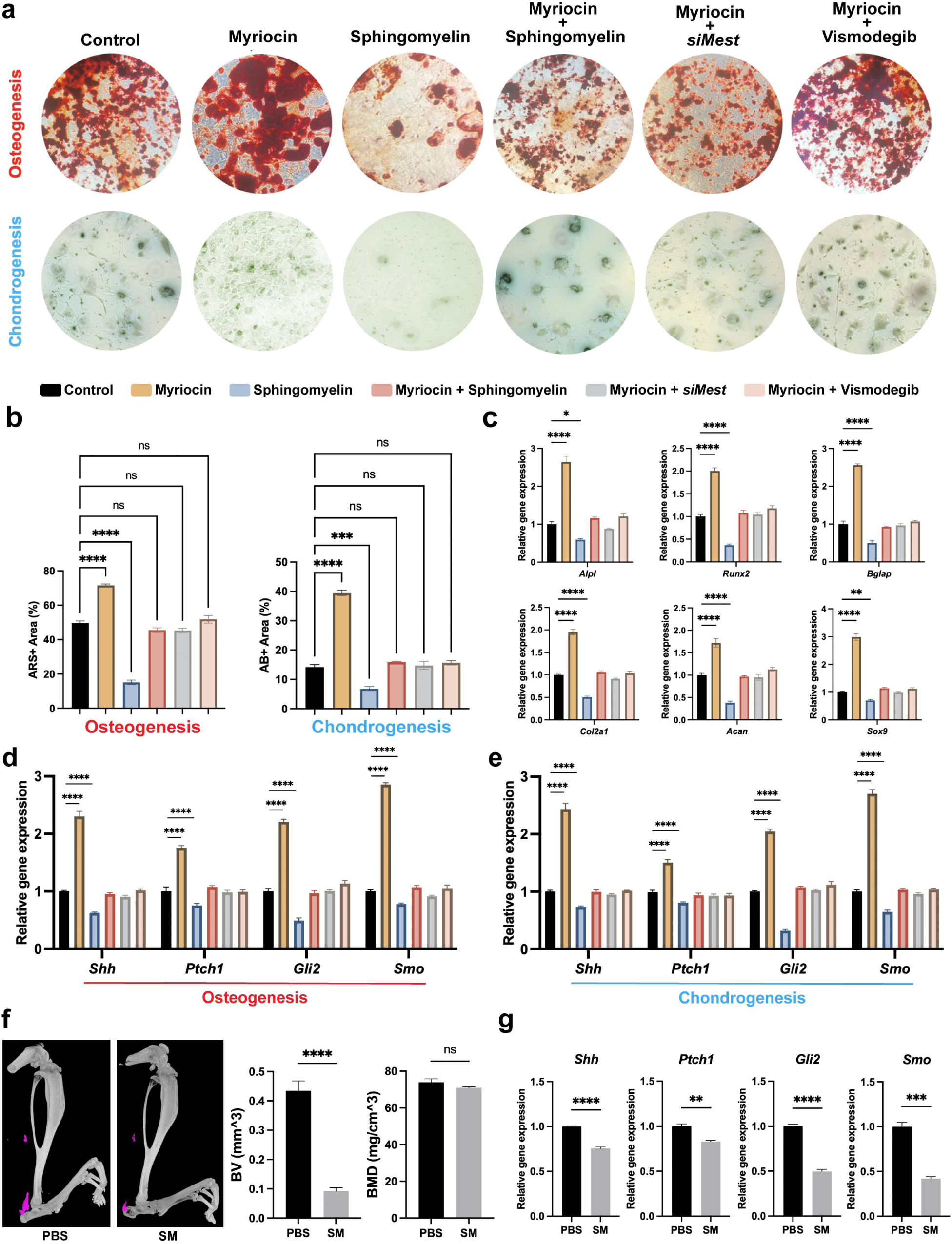
Targeting ciliary sphingomyelin in TSPCs alleviates tHO via Hh signaling. (a) Representative images of alizarin red staining (upper) and alcian blue staining (lower) in TSPCs during in vitro osteogenic (upper) and chondrogenic (lower) differentiation in control, myriocin, sphingomyelin, myriocin + sphingomyelin, myriocin + siMest, and myriocin + vismodegib treated groups. (b) Quantification of the percentages of alizarin red positive (ARS+) and alcian blue positive (AB+) areas in six groups. (c) qPCR showed relative osteogenesis-related genes *(Alpl, Runx2, Bglap*) and chondrogenesis-related genes (*Sox9, Col2a1, Acan*) mRNA expression after in vitro TSPCs osteogenic (upper) or chondrogenic (lower) differentiation in six groups. (d) qPCR showed relative Hh signaling genes (*Shh, Ptch1, Gli2, Smo*) mRNA expression after in vitro TSPCs osteogenic differentiation in six groups. (e) qPCR showed relative Hh signaling genes (*Shh, Ptch1, Gli2, Smo*) mRNA expression after in vitro TSPCs chondrogenic differentiation in six groups. (f) Micro-CT images show tHO of lower limbs between 1X PBS and sphingomyelin (SM) treated groups. Quantification of tHO degree by bone volume (BV, mm^3) and bone mineral density (BMD, mg/cm^3). Independent experiments (n= 4/group) were performed for measurements. (g) qPCR shows relative *Shh, Ptch1, Gli2,* and *Smo* mRNA expression in tHO mice injured site with PBS or SM treatment. Data are presented as means ± SD of three independent assays. Statistical analyses were performed by Student’s t-test for two-group comparison and one-way ANOVA analyses with Tukey’s post hoc test for multiple comparisons. *P < 0.05; **P < 0.01; ***P < 0.005; ****P < 0.001; ns, P > 0.05.

Notably, a study has demonstrated that depletion of sphingomyelin on ciliary membrane can increase the accessibility of cholesterol, thereby amplifying the transmission of the Hh signaling pathway within the cilia^38^. Therefore, we hypothesize that the results may be associated with the activity of the Hh signaling pathway. Smoothened (SMO), a seven-transmembrane receptor protein, serves as a bridge between Hh ligands and GLI transcription factors in the Hh pathway. Direct targeting of SMO has become a trending area in drug research targeting the Hh signaling pathway, with the SMO inhibitor vismodegib being FDA-approved for the treatment of basal cell carcinoma^39^. When vismodegib was administered to TSPCs treated with myriocin, the ARS or AB staining intensity and area was significantly reduced (Figure 9a, b), and the expression of osteogenic (*Alpl,Runx2,*and *Bglap*) and chondrogenic (*Col2a1,Sox9,*and *Acan*) differentiation-related genes was downregulated, with no significant differences compared to the control group (Figure 9c). qPCR analysis of key Hh signaling molecules (*Shh,Ptch1,and Gli2*) during TSPCs osteogenesis (Figure 9d) or chondrogenesis (Figure 9e) showed that myriocin strongly activated Hh signaling during in vitro TSPCs differentiation, while exogenous sphingomyelin had the opposite effect, inhibiting Hh signaling. In contrast, groups treated with myriocin + sphingomyelin, myriocin + *siMest*, and myriocin + vismodegib exhibited Hh signaling gene expression levels that were not significantly different from the control group (Figure 9d).

We also performed in vivo experiments by local injection of 1 mg/kg exogenous sphingomyelin (SM) or 1X PBS (as the control group) into tHO mice injury site two times a week for 8 weeks, resulting in a significantly reduced volume of ectopic bone in tHO mice after SM treatment, with no significant differences in bone mineral density (Figure 9f). qPCR analyses also showed reduced mRNA expression of Hh signaling molecules *Shh, Ptch1, Gli2*, and *Smo* in *SM* treated tHO mice (Figure 9g). These findings suggest that the depletion of ciliary sphingomyelin may facilitate the osteogenic and chondrogenic differentiation of TSPCs through the activation of the Hh signaling, thereby exacerbating tHO progression. This effect can be attenuated by silencing MEST gene expression or using SMO inhibitor vismodegib, which may represent emerging candidate therapeutic targets for tHO.

### 10. scRNA-seq on injured muscle from Nse-Bmp4 tHO model mice

To further validate our findings and enhance the reliability of the results, we reanalyzed the scRNA-seq data (GSE246445) on another tHO model mice which is performed by cardiotoxin (CTX)-induced injury of tibial muscle in *Nse-Bmp4*mice^40^.

After initial quality control, a total of 45287 cells of injured muscle from Nse-Bmp4 mice were clustered into nine clusters based on the reported marker genes^40^ (Figure 10a). While the top two main cell types were skeletal muscle cells and MSC lineages, their proportion during tHO development showed great changes. Notably, the percentage of MSC lineages increased from 14.5% in uninjured group to 66.8% in Day 7 (post injury) group which showed converse trend compared to the skeletal muscle cells (Figure 10b). Consistent with our findings, *Mest*was mainly expressed in MSC lineages (Figure 10c) and showed an increasing expression in injured muscle of *Nse-Bmp4*mice during tHO development (Figure 10d), especially expressing in 60.41% cells in Day 7 group (mainly in MSC lineages). These results demonstrated a significant role of MEST from MSC lineages in tHO pathogenesis which supported our prior findings.

**Figure 10.**
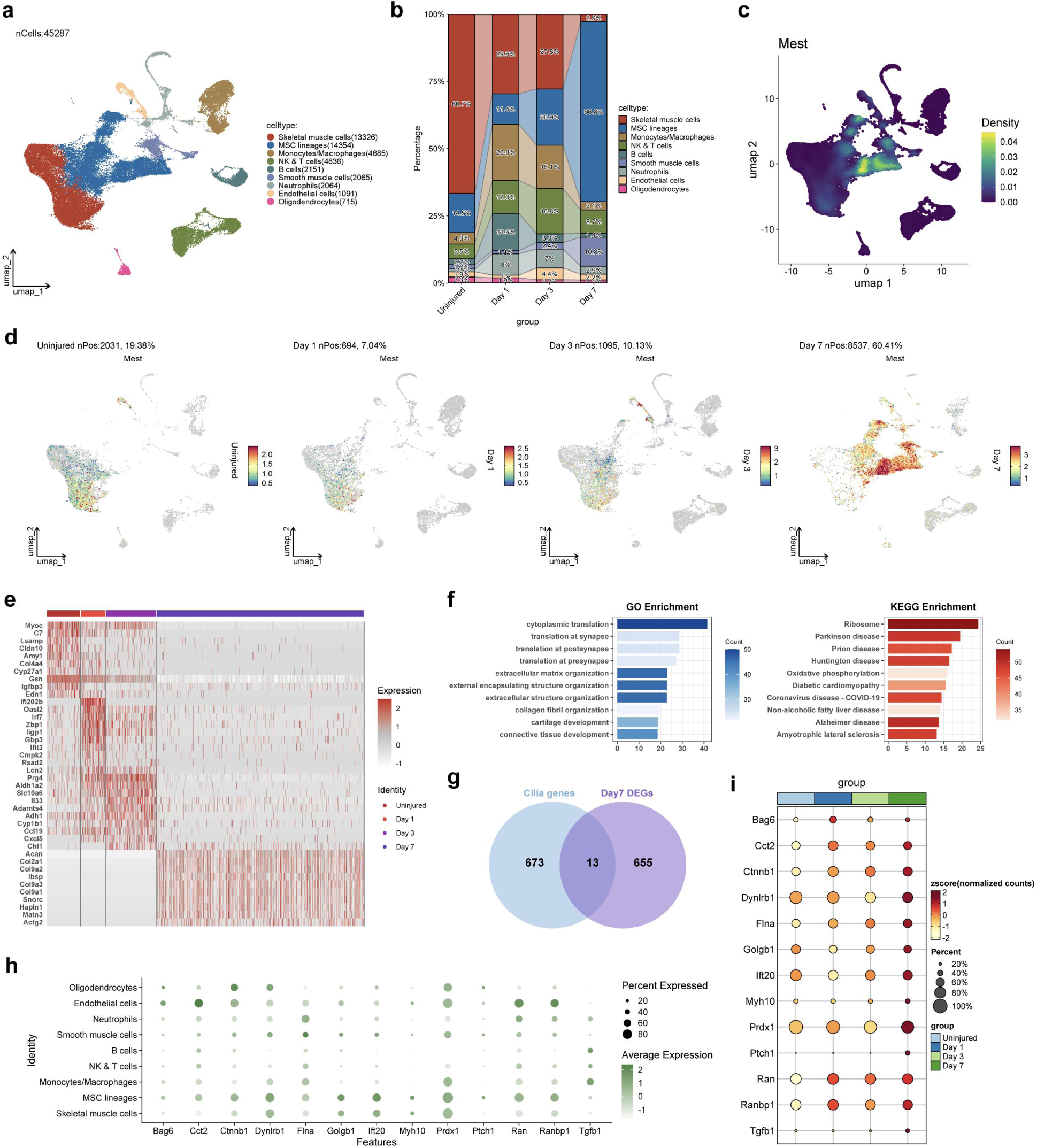
scRNA-seq analysis of the injured tibial muscle from *Nse-Bmp4* tHO mice. (a) UMAP plot of nine clusters identified in 45287 cells in total of *Nse-Bmp4* mice injured muscle. (b) The percentage of each annotated cell type in four groups and its trend. (c) UMAP plot shows the expression density of *Mest* in each cluster. (d) UMAP plot shows the expression of *Mest* from *Nse-Bmp4* mice injured muscle in different groups during tHO development at single-cell levels. (e) Heatmap of the top 10 higher variable genes of four groups in MSC lineages cluster. (f) GO and KEGG enrichment analyses of differential expressed genes (DEGs) in *Nse-Bmp4* mice muscle at 7 days post injury shows top 10 enriched functions and pathways that are important for tHO. (g) Venn plot shows the 13 common genes between cilia genes and DEGs of Day 7 group in MSC lineages cluster. (h) The dot plot visualizes the different expression of 13 common genes in nine clusters. (i) Bubble plot shows the different expression of 13 common genes in four groups of MSC lineages cluster.

Moreover, we analyzed the significant differentiated expressed genes (DEGs) specifically in MSC lineages among four groups (four different timepoints post injury) during tHO development. The top10 higher variable genes in each group were shown by heatmap (Figure 10e). Chondrogenesis-related genes Acan and Col2a1 were top 2 DEGs in Day 7 group, which showed the important role of MSC chondrogenesis in tHO, consistent with our results. GO and KEGG enrichment analysis of DEGs in MSC lineages Day7 group (Figure 10f) also had several common terms with the previous sequencing results shown in Figure 6, indicating that MSC osteochondrogenesis was the key biological process in tHO progression.

Since we have found the potential role of primary cilia and cilia genes in tHO, we were interested in the cilia genes expression in *Nse-Bmp4*tHO model mice at the single-cell level. A total of 686 known cilia genes were obtained from the Syscilia Gold Standard^41^. Meanwhile, we have identified 668 DEGs in Day 7 group of MSC lineages compared with other groups. We then intersected two gene sets and got 13 common genes (Figure 10g), which could be considered as important cilia genes in MSC lineage during tHO development. The expression trends of 13 common cilia genes in each cluster (Figure 10h) or four groups in MSC lineages (Figure 10i) were visualized. Among the 13 genes, IFT20 is the IFT-B complex which also showed increased expression in injured cite of burn/tenotomy tHO model mice as shown in Figure 2 and S2. PTCH1, the key receptor of Hh signaling, was also involved in the 13 common cilia genes, which was consistent with our abovementioned findings. Furthermore, *Sgms2* expression in the MSC lineage cluster declined over time after injury in the Nse-Bmp4 tHO model (Figure S5f), consistent with the results from burn/tenotomy model (Figure 8), implicating ciliary sphingomyelin metabolism (synthesis) in tHO development. Thus, scRNA-seq analysis of the injured muscle from *Nse-Bmp4*tHO mice further demonstrated our conclusions.

## DISCUSSION

The formation of traumatic heterotopic ossification (tHO) is a complex process involving the aberrant differentiation of various types of cells into osteogenic and chondrogenic lineages. Primary cilia are microtubule-based, non-motile organelles that act as cellular antennae, sensing and transducing extracellular signals to regulate various developmental and physiological processes. Several studies have revealed the relationship between primary cilia and skeletal diseases, including defects of skeletal development such as polydactyly^42^ and craniofacial abnormalities^18^. Our previous research found that primary cilia in skeletogenic cells is crucial for BMP and ActA-mediated osteogenesis and chondrogenesis and thereby governs the development of fibrodysplasia ossificans progressiva (FOP)^6^. In this study, we demonstrate that cilia function is significantly upregulated during tHO, with an increased prevalence of ciliated cells in the injury site, particularly among stem/progenitor cells. This observation aligns with previous studies indicating that primary cilia are essential for tissue repair^25^ and regeneration^43^ but extends our understanding by linking cilia activity directly to the pathogenesis of tHO.

Our study reveals that primary cilia positively modulate TSPCs osteochondrogenesis, contributing to the development of tHO. Knockdown of ciliary genes*Ift88*and *Arl3*significantly reduced the osteochondrogenic differentiation capacity of TSPCs and ectopic bone volume in tHO mice. This finding is consistent with the established role of primary cilia in regulating stem/progenitor cell functions. For examples, loss of Numb (a ciliary protein) in spinal neural progenitors lead to reduced cerebellar size via ciliary Hh signaling in mice^44^. The Hh signaling pathway, which is cilia-dependent, is known to be aberrantly activated in postnatal skeletogenesis^45^ and heterotopic ossification in osteoblast^46^. Meanwhile, inactivated PI3K-Akt signaling after ciliary gene IFT80 deficiency results in decreased odontogenic differentiation in dental pulp stem cells^47^. Our study further supports this by showing that ciliary gene knockdown in TSPCs downregulates Hh signaling and inactivates PI3K-Akt signaling, leading to reduced osteochondrogenesis. Other pathway like TGFβ/BMP signaling is also widely studied on both primary cilia^3^ and tHO^48^. Fibrodysplasia ossificans progressiva (FOP) is a genetic form of HO caused by mutations in the BMP type I receptor gene activin A receptor type 1 (ACVR1), while our previous study^6^ demonstrated that primary cilia regulate aberrant osteochondrogenesis via pathological BMP signaling in FOP. Whether ciliary BMP signaling are involved in tHO pathogenesis should be further explored in future research. In summary, we suggested that primary cilia may serve as a crucial regulatory hub, integrating signals that drive the aberrant TSPCs osteochondrogenesis during tHO progression.

A significant contribution of our study is the identification of MEST as a core regulator of tHO for the first time. MEST encodes protein that is widely expressed in mesoderm. As we know, the mesoderm develops into the skin, muscle, bone, and other connective tissues in our body, which indicate the potential role of MEST in skeletogenesis. Our scRNA-seq results showed that MEST is predominantly expressed in stem/progenitor cells and is upregulated during tHO progression. A recent study reported that MEST specifically expressed in mesenchymal cells and induced bone formation during mouse digit tip regeneration^49^. In our study, scRNA-seq showed that the expression levels of osteogenic and chondrogenic genes in Mest highly expressed stem/progenitor cells were higher than Mest low expressed stem/progenitor cells. Moreover, knockdown of MEST in TSPCs attenuated their osteogenic and chondrogenic differentiation capacities, while *shMest*significantly alleviate severity of tHO in mice, indicating a pivotal role of MEST in promoting tHO by regulating TSPCs differentiation. Interestingly, MEST can regulate the cilia length but not counts in TSPCs though it has no co-localization with primary cilia. However, the specific mechanism of how MEST act on cilia remains unknown. Additional studies are needed to characterize how MEST modulates ciliary structural remodeling and interacts with key ciliary functional components. Overexpressing MEST in *shIft88*TSPCs can rescue TSPCs reduced osteochondrogenic abilities, suggesting the effects of primary cilia on TSPCs osteochondrogenesis and tHO is dependent on MEST activity. This finding is innovative as it uncovers a novel regulatory axis involving cilia and MEST, highlighting a potential therapeutic target for tHO.

Our study elucidates the molecular mechanisms through which MEST regulates tHO. We showed that silencing MEST can suppress Hh and PI3K-Akt signaling and attenuate TSPCs osteochondrogenesis in vitro. Overexpression of MEST promote TSPCs osteochondrogenesis via activating Hh and PI3K-Akt signaling, and the effect can be rescued by Hh or PI3K-Akt inhibitor. Furthermore, we show that the GLI2 transcription factor, a key activator of the Hh pathway, binds to and promotes MEST transcription. This finding is supported by previous studies indicating that GLI2 is a master regulator of bone homeostasis^50^ and that its activity is tightly regulated by cilia function^51^. Taken together, this study indicated that GLI2 transcriptional factor activates MEST transcription and increase its gene expression, which in turn enhances TSPCs osteogenesis and tHO progression via Hh and PI3K-Akt signaling.

In the presence of external stimuli, the selective influx and efflux of specific lipids in primary cilia can dynamically modulate cell functions^13^. This process is intricately linked to adipogenesis^34^ and diseases such as polycystic kidney disease^17^. Thus, another novel aspect of our study is the investigation of ciliary lipid metabolism in TSPCs during tHO. By untargeted lipidomic after isolation of primary cilia, we show that MEST regulates the abundance of various important lipids within TSPCs cilia, especially with sphingomyelin levels being significantly upregulated upon MEST knockdown. This finding is intriguing as sphingomyelins are known to interact with cholesterol to regulate ciliary signaling processes^38^. Recent studies showed that SGMS2 mediated SM synthesis is involved in pancreatic cancer^52^ and atherosclerosis^53^. Meanwhile, it is worth noting that SGMS2 is associated with early-onset osteoporosis^54^ and skeletal dysplasia^36^, indicating its potential role in bone diseases. Our findings first indicated that SGMS2 expression and SM synthesis may be activated by MEST knockdown, resulting in the increased ciliary SM levels in *siMest*treated TSPCs, which may provide novel insights into the treatment of tHO. Further studies should further concentrate on the mechanism by which MEST may regulate SM metabolism in cilia. To explore the translational value of targeted SM metabolism in tHO, we demonstrated that exogenous sphingomyelin treatment inhibits TSPCs osteochondrogenesis and reduce ectopic bone volume in tHO mice. This effect is mediated through the ciliary Hh signaling, as evidenced by the activation of Hh signaling upon myriocin treatment and its inhibition by exogenous sphingomyelin. Vismodegib, a SMO inhibitor targeting Hh signaling, can also diminish osteogenesis and chondrogenesis of TSPCs. The levels of other sphingolipids and glycerophospholipids were also partly affected in *siMest*treated TSPCs, indicating that other ciliary lipids except SM can also be involved in tHO pathogenesis and need further studies to validate. These findings provide a new perspective on the role of lipid metabolism in tHO and suggest that targeting ciliary sphingomyelin levels may represent a potential therapeutic strategy.

Several limitations should be acknowledged in the study. First, our study primarily focuses on the role of primary cilia in TSPCs, while other cell types^55,56^ involved in tHO (such as endothelial cells, muscle satellite cells, mesenchymal stem cells, adipose-derived stem cells, fibroblasts, and immune cells) were not extensively investigated. Future studies should explore the cilia-dependent functions of these cells and their interactions with TSPCs during tHO. Second, our in vivo experiments were limited to local injection of AAV vectors carrying *shRNA*, which may not fully recapitulate the complex physiological environment of tHO. Further studies using genetically modified animal models are needed to validate our findings. Third, while we identified MEST as a key regulator of tHO, the detailed molecular mechanisms through which MEST modulates sphingomyelin metabolism within cilia remain unclear. Future studies should investigate the specific interactions between MEST and lipid metabolic enzymes and their impact on cilia function as the regulator of tHO. Finally, we did not explore whether and how ECM stiffness alters cilia and their downstream signaling in tHO though 1) mechanical stimuli can regulate bone formation via ciliary mechanical transduction^57^; 2) results of bioinformatic analyses (Figure 5) show ECM-related biological processes may play a crucial role in TSPCs osteochondrogenesis or tHO formation. Future studies should clarify how ECM stiffness after injury influence local ectopic bone formation through ciliary mechanotransduction in tHO.

The schematic diagram of the study is shown in Figure 11. Our study opens several avenues for future research. First, since primary cilia also located in other cell types during tHO development, further investigation into the role of primary cilia in other cell types involved in tHO is warranted. This will provide a more comprehensive understanding of the cilia-mediated molecular mechanisms underlying tHO progression. Second, exploring the therapeutic potential of targeting cilia function or MEST expression in preclinical models of tHO is an important next step. This could involve the development of small molecules or gene therapies aimed at modulating cilia activity or inhibiting MEST function. Third, elucidating the detailed molecular mechanisms through which MEST regulates sphingomyelin or other lipid metabolism within cilia will provide new insights into the pathogenesis of tHO and identify potential therapeutic targets. Finally, translating these findings into clinical applications will require rigorous preclinical and clinical trials to assess the safety and efficacy of cilia-targeted therapies for tHO.

**Figure 11.**
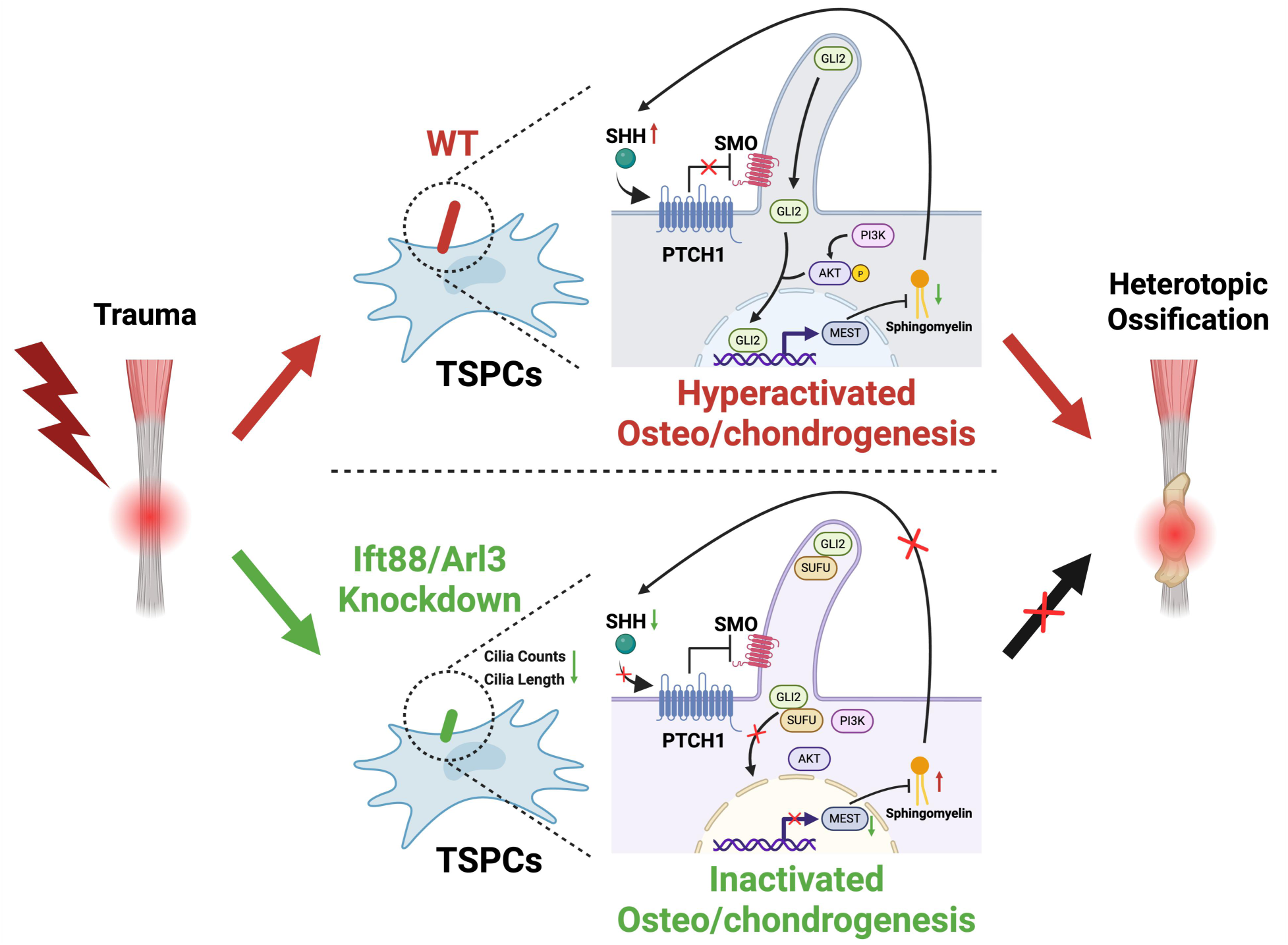
Schematic diagram of the study. SHH is unregulated within the primary cilia of TSPCs after trauma, activating the Hh signaling through its interaction with PTCH1 and SMO receptors. Subsequently, the GLI2 transcription factor activated and translocated to the nucleus, where it promotes the expression of the MEST gene. Concurrently, the phosphorylation levels of PI3K and AKT are enhanced. The upregulation of MEST expression inhibits sphingomyelin synthesis and leads to a reduction in sphingomyelin levels within the cilia, further amplifying the activation of the ciliary Hh signaling. This positive feedback loop dramatically enhances the osteogenic and chondrogenic differentiation potential of TSPCs, resulting in localized ectopic bone deposition and tHO. Conversely, when the ciliary genes Ift88 and Arl3 are knocked down in TSPCs, the production of SHH ligands is markedly reduced. Thus, SMO is inhibited by PTCH1, preventing the activation of GLI2. Additionally, the phosphorylation of PI3K and AKT is suppressed. The reduction in MEST expression subsequently increases the sphingomyelin within the cilia, which further inhibits the ciliary Hh signaling. As a result, the osteogenic and chondrogenic differentiation capacity of TSPCs is significantly diminished, thereby alleviating the occurrence and development of tHO.

## MATERIALS AND METHODS

### 1. Experimental tHO mice

Male C57/BL6 mice weighing approximately 20 g and aged 6–8 weeks bought from Hangzhou Ziyuan Laboratory Animal Technology Co., Ltd were used for construction of burn/tenotomy tHO mice model as previously reported^21^. Briefly, mice were anesthetized via intraperitoneal injection of sodium pentobarbital. Hair was removed from the left hind limb (from the knee to the ankle) and a 2×3 cm area on the left side of the midline of the spine using an electric clipper for surgical preparation. A longitudinal incision was made along the Achilles tendon to expose the gastrocnemius tendon, which was then transected transversely. The skin was sutured using 5-0 surgical thread. Subsequently, a 2×3×3 cm aluminum block preheated to 60°C in a water bath was applied to the 2×3 cm area on the left side of the midline of the spine for approximately 17 seconds without additional pressure. The surgical site was disinfected with an alcohol swab postoperatively, and mice were housed individually. Antibiotics and analgesics were administered as needed. The success of the tHO model was evaluated 8–10 weeks post-surgery. Local injection of 1 mg/kg sphingomyelin (#HY-113498, MCE) and 1X PBS at the same dose into the injured muscle in tHO mice were performed two times per week.

### 2. Micro-CT

After euthanasia, the lower leg and foot of the mice were harvested and fixed overnight at 4°C in 10% formalin solution. Subsequently, the samples were subjected to high-resolution micro-CT using a SkyScan CT reconstruction software (Bruker Corporation). Three-dimensional modeling and additional data analysis were performed using 3D.SUITE (Bruker Corporation).

The entire ectopic bone region of the specimen was selected as the region of interest (ROI) for quantification of bone volume (BV). Additionally, the distal tibial epiphysis of the mouse was selected as the ROI for quantification of bone mineral density (BMD) and trabecular thickness (Tb.Th).

### 3. Immunofluorescence

For immunofluorescence staining of tissues and cells, samples were prepared by fixing in 4% paraformaldehyde at 4°C for 24 hours (tissues) or at room temperature for 15 minutes (cells). Tissues were subsequently deparaffinized and rehydrated through gradient ethanol, while cells were permeabilized with ice-cold methanol or 0.3% Triton X-100 (Solarbio Life Sciences) as needed. Both tissues and cells were blocked with 10% normal goat serum or 1% BSA to reduce nonspecific binding. Primary antibodies, raised in different host species, were applied and incubated overnight at 4°C. After washing, fluorophore-conjugated secondary antibodies (#A-11034, Thermo Fisher Scientific) were added and incubated for 1 hour at room temperature in the dark. Nuclei were stained with DAPI for 10–15 minutes, followed by washing. Sections and cells were mounted with an anti-fade medium and imaged using Nikon ECLIPSE Ni microscopic system. Primary antibodies used in immunofluorescence are listed in Table S3.

### 4. Histology and immunohistochemistry

Following euthanasia, the tHO mice tendon tissues were collected and fixed in 4% paraformaldehyde at 4°C for 24 hours. After three washes with PBS, the samples were decalcified in 10% EDTA at 4°C for 21 days. The tissues were then embedded in paraffin, and sagittal sections of the Achilles tendon were obtained via paraffin microtomy. Histological staining was performed using hematoxylin and eosin (H&E), Masson’s trichrome, and Von Kossa staining to evaluate ectopic ossification in the Achilles tendon. The sections were dehydrated in ethanol, mounted, and imaged using Nikon ECLIPSE Ni microscopic system.

For immunohistochemistry on paraffin-embedded sections, the slides were first deparaffinized in xylene and rehydrated through a graded ethanol series. Endogenous peroxidase activity was quenched with 3% hydrogen peroxide in methanol. Antigen retrieval was performed by heating the sections in citrate buffer (pH 6.0) for 10 minutes at 95–100°C. Sections were then blocked with 10% fetal bovine serum or 1% BSA for 1 hour at room temperature. Primary antibodies were applied and incubated either for 1 hour at room temperature or overnight at 4°C, followed by washing with PBS. Secondary antibodies (#Ab205718, abcam) were added and incubated for 30 minutes at room temperature. For visualization, sections were developed with DAB substrate and counterstained with hematoxylin. Finally, the sections were dehydrated, cleared, and mounted with neutral gum for microscopic examination using Nikon ECLIPSE Ni microscopic system. Primary antibodies used in immunohistochemistry are listed in Table S3.

### 5. Isolation of TSPCs and cell culture

The isolation of TSPCs was conducted as previously reported^58^. Briefly, 6- to 8-week-old C57/BL6 mice were sacrificed, and their entire gastrocnemius tendons were aseptically harvested. The tendons were washed three times with 1X PBS and finely minced using ophthalmic scissors. The minced tissue was collected in a 15 ml centrifuge tube and digested with 3 ml of 3 mg/ml collagenase type I (#SCR103, Sigma-Aldrich) at 37°C in a shaking incubator for 45 minutes. The mixture was then centrifuged at 500 rpm for 5 minutes to remove the collagenase. Subsequently, the pellet was resuspended in 3 ml of 2 U/ml dispase type II (#4942078001, Roche) and digested at 37°C for another 45 minutes. After centrifugation at 500 rpm for 1 minute to remove the dispase, the cell pellet was resuspended in 3 ml of low-glucose DMEM supplemented with 10% fetal bovine serum (FBS) and 1% penicillin/streptomycin. The cell suspension was filtered through a 70 µm cell strainer into a 10 cm culture dish and supplemented with 6-8 ml of complete culture medium. The medium was changed for the first time after 3 days and then every other day. When the cells reached 80% confluence (7-10 days later), they were passaged for further experimental use. The drugs or substances added to TSPCs and the concentration in the cell experiment are listed below: 10 μM myriocin (#HY-N6798, MCE), 20 μM sphingomyelin (#HY-113498, MCE), 10 μM vismodegib (#HY-10440, MCE), 4 μM GANT61 (#HY-13901, MCE), 20 μM LY294002 (#HY-10108).

### 6. Flow cytometry

Fluorescence-activated cell sorting (FACS) analysis of mouse TSPCs were performed following the protocols of previous studies^31,59^. We first removed the red blood cells from the TSPCs using RBC Lysis Buffer (#46232, CST) and resuspend the cells in FACS buffer (1X PBS + 2% FBS + 1mM EDTA). The extracted TSPCs were stained with antibodies for 30 min at 4°C in the dark. Cells were then washed twice with FACS buffer to remove unbound antibodies. Flow cytometry was performed using BD FACSAria^TM^ III. Data were analyzed using BD FACS Diva software and FlowJo software.

### 7. Plasmid construction and transfection

The pLVX-shRNA-puro lentivirus vector carrying shRNA was constructed by Shanghai OBiO Technology Inc. for targeted gene knockdown in vitro. For lentivirus-mediated transduction in TSPCs, 8×10⁴ cells were seeded into each well of a 6-well plate. When the cell density reached 20%–30%, cells were transduced with lentivirus in the presence of polybrene (8– 10 hours). Subsequently, stable transduced cells were selected using 2 µg/ml puromycin for 5–10 days. The successful establishment of stable cell lines was verified by quantitative qPCR and Western blot (Figure S7a).

The pAAV-U6-shRNA-CMV-MCS-WPRE adeno-associated virus (AAV) vector carrying shRNA was constructed by Shanghai OBiO Technology Inc. for targeted gene knockdown in vivo. The shRNA sequences used are listed in Table S4. Forty-eight hours after tHO induction in mice, 10 µL of the AAV was injected intramuscularly into the medial side of the injured hind limb. Subsequent injections were administered every 3–5 days to achieve sustained knockdown of the specific gene expression. The successful establishment of gene knockdown was verified by quantitative qPCR and Western blot (Figure S7b, c).

The pCDNA3.1 plasmid carrying the full-length cDNA of Gli2 or Mest was constructed for targeted gene overexpression in vitro. When cells reached 50%-60% confluence, plasmid (Gli2 pcDNA3.1 plasmid, Mest pcDNA3.1 plasmid, or pcDNA3.1 control plasmid) transfection was carried out using Lipofectamine 3000 (#L3000001, Invitrogen) according to the manufacturer’s protocol. The cell culture medium was refreshed 24h after transfection.

### 8. siRNA transfection

The siRNA sequences used are listed in Table S4. For siRNA transfection, 1×10^5^ cells were seeded into each well of a 6-well plate. When the cell density reached 50%, cells were transfected with siRNA using Lipofectamine RNAiMAX (Invitrogen). Specifically, the siRNA was mixed with Lipofectamine RNAiMAX and incubated at room temperature for 10 minutes. The mixture was then added to cells cultured in Opti-MEM reduced-serum medium. After incubation for 48–72 hours, cells were harvested for subsequent experiments.

### 9. Induction of TSPCs osteogenic and chondrogenic differentiation

To induce osteogenic differentiation of TSPCs, gelatin-coated 6-well plates were used to promote cell attachment and growth. TSPCs were seeded at a density of 1×10^4^ cells/cm^2^ and cultured in osteogenic induction medium containing 10 mM β-glycerophosphate, 50 µg/ml ascorbic acid, and 100 nM dexamethasone. The medium was changed every 2-3 days to maintain optimal conditions. Osteogenic differentiation was assessed by alizarin red staining after 2-3 weeks to visualize calcium deposits. Additionally, qPCR was performed to evaluate the expression of osteogenesis related genes.

To induce chondrogenic differentiation of TSPCs, cells were seeded at a density of 1×10^4^ cells/cm^2^ on gelatin-coated 6-well plates and cultured in chondrogenic induction medium containing 10 ng/ml transforming growth factor-β3 (TGF-β3), 100 nM dexamethasone, 50 µg/ml ascorbic acid, and 10 mM β-glycerophosphate. The medium was refreshed every 2-3 days to ensure optimal conditions for differentiation. Chondrogenic differentiation was assessed by histological staining with alcian blue to detect glycosaminoglycans at 2-3 weeks. qPCR was also performed to evaluate the expression of chondrogenic markers.

### 10. Quantitative polymerase chain reaction (qPCR)

To assess the mRNA levels of the target genes, total RNA was purified from TSPCs using an RNA Easy Fast Tissue/Cell Kit (#DP451, TIANGEN). The extracted mRNA was reverse-transcribed into cDNA using FastKing gDNA Dispelling RT SuperMix (#KR118, TIANGEN). Quantitative PCR was performed using the ABI Step One Plus system (Applied Biosystems). Each reaction mixture (final volume of 20 µL) contained 1 µL of total cDNA, Power SYBR Green PCR Master Mix (#A57155, Invitrogen), and gene-specific primers. The PCR protocol included an initial denaturation at 95°C for 10 minutes, followed by 45 cycles of 95°C for 20 seconds, 72°C for 30 seconds, and a final dissociation curve analysis from 60°C to 95°C at a heating rate of 0.1°C per second. GAPDH was used as an internal control. Relative gene expression was calculated using the 2^−ΔΔCt^ method. Primers used in the study are listed in Table S5.

### 11. Western blot

To perform western blot analysis, proteins were extracted from cells by washing with PBS, lysing in RIPA buffer (#R0278, Sigma-Aldrich) containing Protease & Phosphatase Inhibitor Cocktail (#78441, Thermo Fisher Scientific), and sonicating on ice. The lysates were centrifuged at 4°C to collect the supernatant, and protein concentrations were determined using a BCA Protein Assay Kit (#P0011, Beyotime). Equal amounts of protein were denatured by heating in SDS loading buffer, separated by SDS-PAGE, and transferred to PVDF or nitrocellulose membranes. Membranes were blocked with 5% skim milk in TBST to reduce nonspecific binding. Primary antibodies were applied overnight at 4°C, followed by washing and incubation with secondary antibodies conjugated to horseradish peroxidase (#SA00001-2, Proteintech Inc.) for 1 hour at room temperature. After additional washing, the membranes were developed using Super ECL Detection Reagent (#36208ES60, YEASEN) and imaged with ChemiDoc XRS+ Gel Imaging System (Bio-Rad). Band intensities were quantified using ImageJ software, and target protein expression levels were normalized to those of the internal control. Primary antibodies used in western blot are listed in Table S3.

### 12. RNA-seq

Total RNA was isolated from samples using RNA Easy Fast Tissue/Cell Kit (#DP451, TIANGEN). The purity and concentration of the extracted RNA were assessed using the NanoDrop One spectrophotometer (Thermo Fisher Scientific). RNA integrity was further evaluated using the Agilent 2100 Bioanalyzer (Agilent Technologies). Subsequently, RNA sequencing libraries were prepared using the VAHTS Universal V6 RNA-seq Library Prep Kit (Vazyme). The transcriptome sequencing and subsequent bioinformatics analysis were performed by Shanghai OE Biotech. Sequencing libraries were sequenced on the Illumina Novaseq 6000 platform to generate 150-bp paired-end reads. The raw sequencing reads in FASTQ format were processed using the fastp software to remove low-quality reads, yielding clean reads for subsequent data analysis. The clean reads were aligned to the reference genome using the HISAT2 software, and gene expression levels were calculated in terms of fragments per kilobase of transcript per million mapped reads (FPKM). Additionally, read counts for each gene were obtained using HTSeq-count. Subsequent analyses, including principal component analysis (PCA), differential gene expression analysis, and enrichment analysis, were performed using the R software.

### 13. Untargeted lipidomics using liquid chromatography-tandem mass spectrometry (LC-MS/MS)

TSPCs were cultured in serum-free medium to promote cilium growth until confluence. The cells were washed with PBS and subjected to shear force by rotating at 360 rpm in a shaking incubator at 37°C for 4 minutes to detach cilia as previously reported^35^. The cell debris was removed by centrifugation at 1000×g for 10 minutes at 4°C, followed by ultracentrifugation at 40,000×g for 30 minutes at 4°C to isolate the ciliary suspension from the supernatant. The ciliary suspension was then used for untargeted lipidomics sequencing.

For lipid extraction, 200 µL of pre-chilled methanol/water (3:1, v/v) was added to the ciliary suspension, followed by sonication on ice for 15 minutes. Then, 1 mL of pre-chilled methyl tert-butyl ether (MTBE) was added and vortexed thoroughly. The mixture was rotated at 4°C for 1 hour, followed by sonication on ice for another 30 minutes. After adding 100 µL of water and vortexing for 1 minute, the mixture was left at room temperature for 10 minutes. The sample was then centrifuged at 14,000×g for 15 minutes at 4°C. The supernatant was collected, and the pellet was dried under a stream of nitrogen. The dried pellet was re-dissolved in 200 µL of SDT buffer. Protein concentration was measured using the BCA assay. Based on the protein content, the cell quantity in each sample was estimated, and equal cell amounts were taken from each supernatant. The supernatant was dried using a high-speed vacuum concentrator, re-dissolved in 60 µL of pre-chilled isopropanol/methanol (1:1, v/v), and centrifuged at 20,000×g for 20 minutes at 4°C. The supernatant was vortexed and transferred to an injection vial for analysis. The entire analysis was performed at 4°C using an automatic sampler. The samples were analyzed using a SHIMADZU-LC30 ultra-high-performance liquid chromatography (UHPLC) system equipped with an ACQUITY UPLC® HSS C18 column (2.1×100 mm, 1.9 µm) (Waters, Milford, MA, USA). To avoid signal drift, samples were analyzed in a random order. A quality control (QC) sample was inserted every 5-10 experimental samples to monitor system stability and data reliability. The samples were analyzed in both positive and negative ion modes using electrospray ionization (ESI). After separation by UHPLC, the samples were analyzed using a QExactive Plus mass spectrometer (Thermo Fisher Scientific). Lipid identification and quantification were performed using the MSDIAL software (Version 5.2.240218.2-net472) with the following parameters: precursor mass tolerance of 10 ppm, product mass tolerance of 10 ppm, retention time alignment of 0.1 min, and peak area filtering to reduce false positives. Lipids with relative standard deviations (RSD) exceeding 30% in QC samples were excluded. The data were normalized to the total peak area, preprocessed with UV-scaling, and subjected to multivariate statistical analyses, including unsupervised principal component analysis (PCA), supervised partial least squares discriminant analysis (PLS-DA), and orthogonal partial least squares discriminant analysis (OPLS-DA). Univariate statistical analyses included Student’s t-test and fold-change analysis, and volcano plots and hierarchical clustering were generated using R software.

### 14. Dual-luciferase reporter assay

The Dual-Luciferase Reporter Assay was performed using the Dual-Luciferase Reporter Assay System (Promega). The firefly luciferase reporter gene was cloned downstream of the wild-type (WT) or mutant (MUT) pGL4.10-Mest promoter regions and co-transfected into TSPCs with pcDNA3.1-GLI2 and Renilla expression vector pRL-TK. Following a 48-hour incubation, cell lysates were prepared, and firefly luciferase activity was quantified and normalized to Renilla luciferase activity as an internal control.

### 15. Chromatin immunoprecipitation (ChIP) assay

The ChIP assay was performed according to the previous study^60^ using Pierce^TM^ Agarose ChIP Kit (#26156, Thermo Fisher Scientific). In brief, TSPCs were fixed with 1% formaldehyde, quenched with 1X glycine solution, and lysed using Lysis Buffer supplemented with protease inhibitors. Chromatin was digested with ChIP-grade micrococcal nuclease. The digested chromatin was incubated overnight at 4°C with GLI2 antibody in Dilution Buffer. ChIP-grade Protein A/G Plus Agarose beads were added to each immunoprecipitation (IP), followed by centrifugation and washing with Wash Buffer. Elution was performed using 150 μL IP Elution Buffer, 6 μL 5 M NaCl, and 2 μL 20 mg/mL Proteinase K. DNA fragments were analyzed by real-time PCR. The primer sequences for ChIP-qPCR were designed based on +/-150 bp of the two predicted sites on MEST promoter region and provided in Table S5.

### 16. Bioinformatics analysis

The scRNA sequencing dataset from the GEO database (GSE126060 and GSE246445). Dimensionality reduction, clustering, and cell-type identification of the scRNA sequencing data were performed using the Seurat package in R software. Additionally, bulk RNA sequencing data from the heterotopic ossification sites of mice three weeks post-trauma was also obtained from the GEO database (GSE233201). Differential gene expression analysis was conducted using the DESeq2 package, while enrichment analysis was performed with the clusterProfiler package. Transcription factor binding sites were predicted using the JASPAR database (https://jaspar.elixir.no/).

### 17. Statistical analysis

All statistical analyses were performed using SPSS software and R Studio. Data are presented as mean ± standard deviation (SD) of three independent assays. Statistical analyses were performed by Student’s t-test for two-group comparison and one-way ANOVA analyses with Tukey’s post hoc test for multiple comparisons. Statistical significance was set at P < 0.05.

## List of Supplementary Materials

Figure S1 to S7 for multiple supplementary figures

Table S1 to S5 for multiple supplementary tables (including Excel files)

## Acknowledgments

We thank Airu Zhu at Shanghai Bioprofile Technology Company. Ltd for her technical support in mass spectroscopy. Schematic diagrams (Figure 1a, 6a, 6b, and S7) were created with BioRender (https://www.biorender.com/).

## Author contributions

Conceptualization: HJ, XHZ; Methodology: BWL, JQZ; Investigation: BWL, YG; Visualization: BWL, ZLS; Funding acquisition: HJ, RG, XHZ; Project administration: HJ, XHZ; Supervision: JQZ, RG; Writing – original draft: BWL, YG; Writing – review & editing: HJ, XHZ

## Data and materials availability

The data that support the findings of this study are available from corresponding author upon reasonable request. Publicly available datasets were analyzed in this study, these can be found in Gene Expression Omnibus (GSE126060, GSE246445 and GSE233201) of National Center for Biotechnology Information.

## Ethical approval

All animal procedures were conducted in accordance with the guidelines of the Ethics Committee on Animal Experiments of the Naval Medical University.

