## Supplementary Materials for "MEST inhibits ciliary sphingomyelin synthesis to promote tendon stem/progenitor cells osteochondrogenesis in traumatic heterotopic ossification"

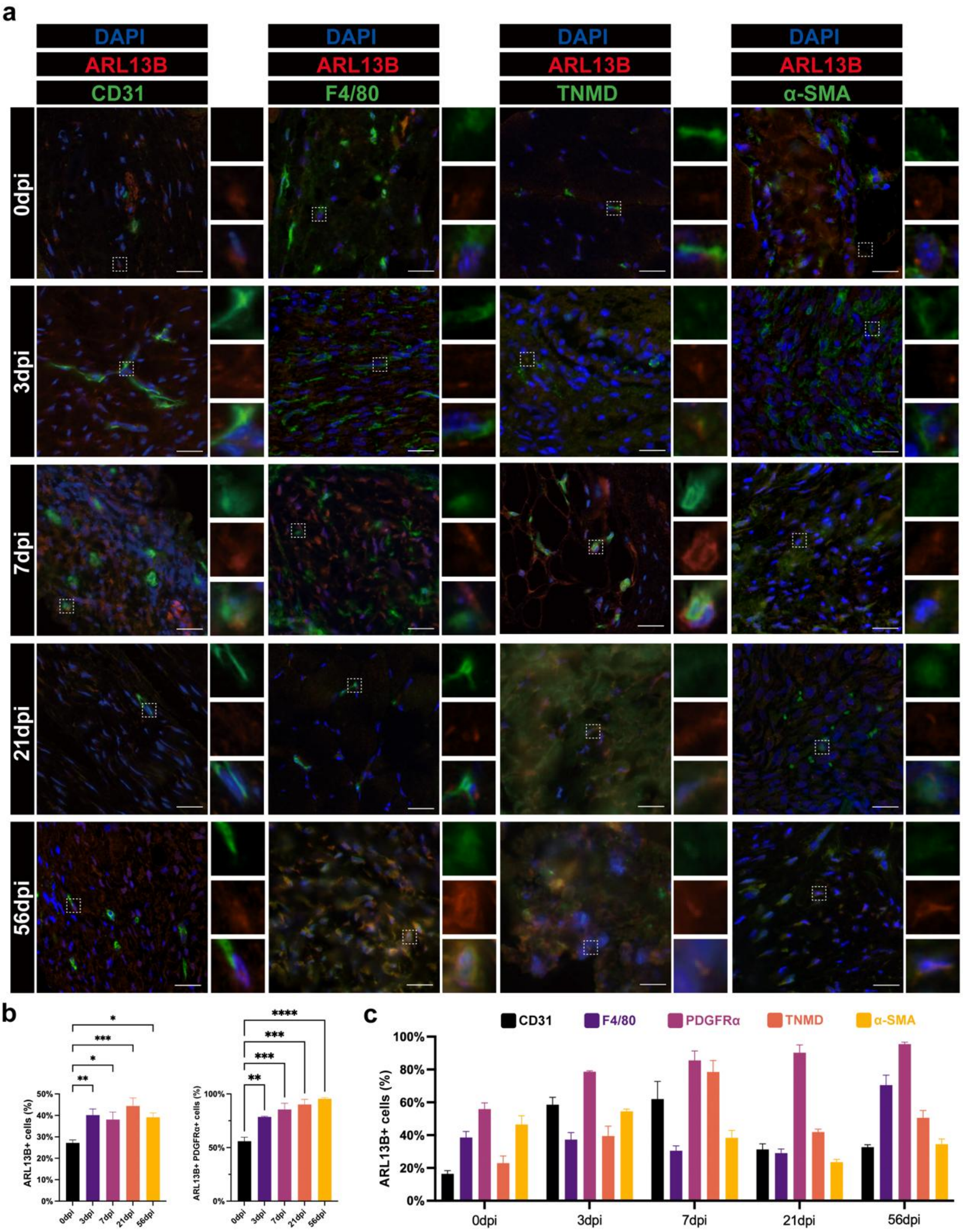


**Figure S1.** (a) Immunofluorescence images of different cell type markers (CD31, F4/80, TNMD, α-SMA) (green) and ARL13B (red) at different time points during tHO development. DAPI staining indicates nuclei (blue), scale bar 30 μm. (b) The percentage of ARL13B+ cells at different time points in all cells (left) and PDGFRα+ cells (right) after injury are shown (n= 3). (c) The percentage of ARL13+ cells in five types of cells (CD31+, F4/80+, PDGFRα+, TNMD+, and α-SMA+ cells) are shown (n= 3). Data are presented as means ± SD of three independent assays.

Statistical analyses were performed by one-way ANOVA analyses with Tukey's post hoc test for multiple comparisons. *$P < 0.05$; **$P < 0.01$; ***$P < 0.005$; ****$P < 0.001$.

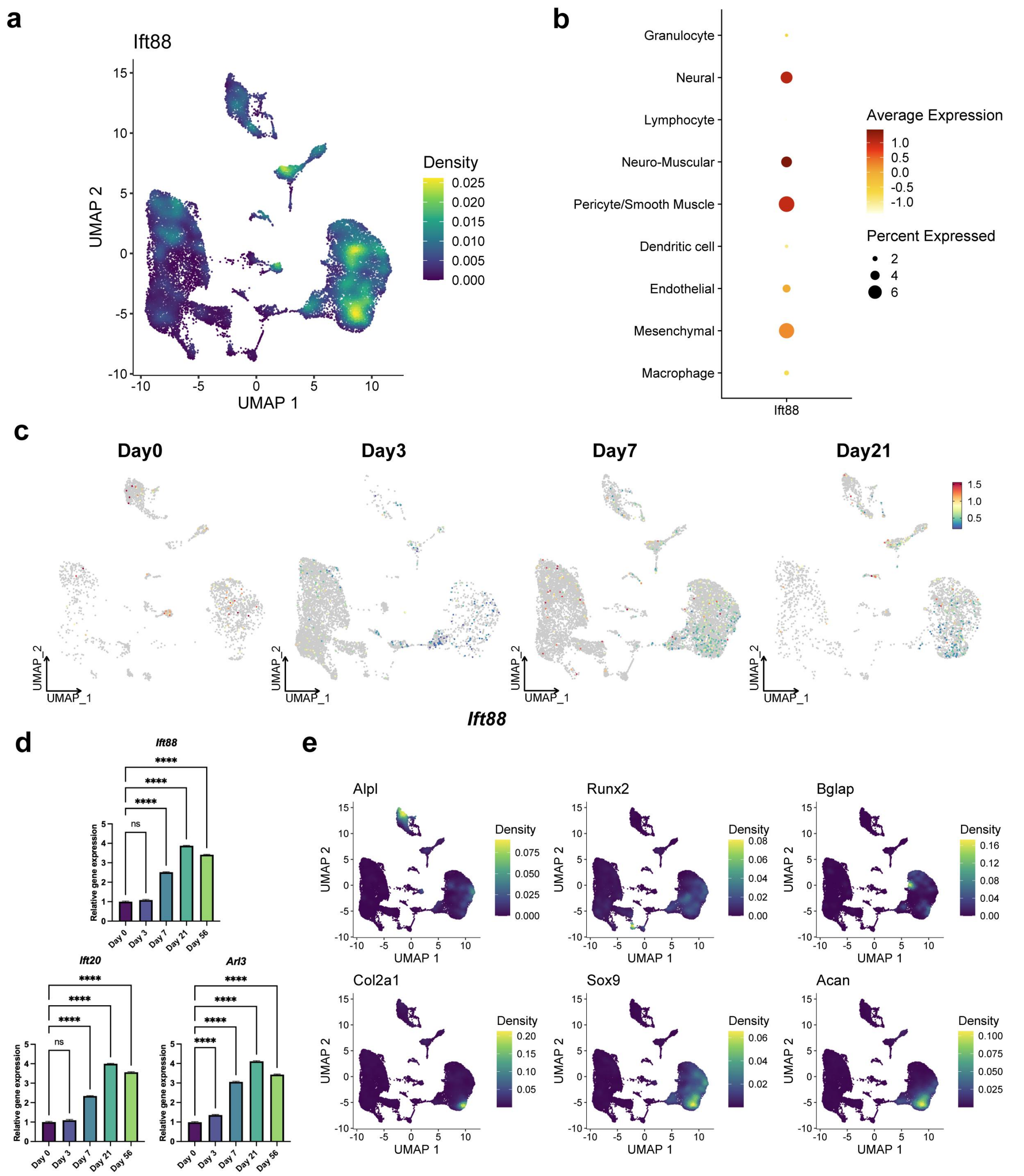


**Figure S2.** (a) The expression density of *Ift88* in each cluster are shown by UMAP plot. (b) The dot plot visualizes the different expression of *Ift88* in nine clusters. (c) UMAP plot shows the expression of *Ift88* at different time points (Day 0 ,3, 7, 21) during tHO development. (d) qPCR showed ciliary genes *Ift88, Ift20*, and *Arl3* mRNA expression in injured tendon tissues from tHO mice at different time points (Day 0, 3, 7, 21). (e) The expression density of osteogenesis-related (*Alpl, Runx2*, and *Bglap*) and chondrogenesis-related (*Col2a1, Sox9,* and *Acan*) genes in each cluster are shown by UMAP plot. Data are presented as means ± SD of three independent assays. Statistical analyses were performed by one-way ANOVA analyses with Tukey's post hoc test for multiple comparisons. ***P < 0.005.

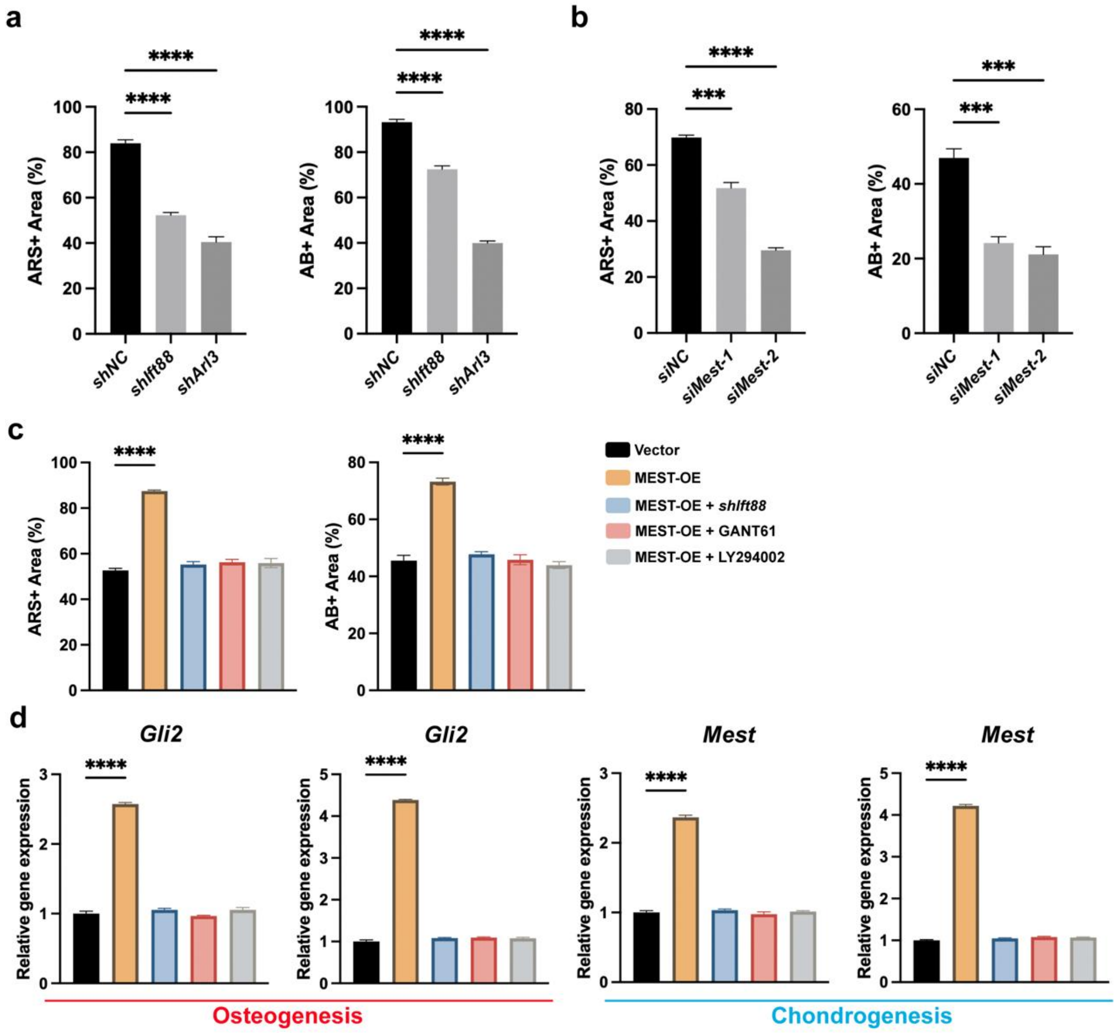


**Figure S3. (a)** Quantification of the percentages of (alizarin red) ARS+ and alcian blue (AB+) areas in *shNC, shIft88*, and *shArl3* groups. (b) Quantification of the percentages of ARS+ and AB+ areas in *siNC, siMest-1, and siMest-2* groups. (c) Quantification of the percentages of ARS+ and AB+ areas in *Vector, MEST-OE, MEST-OE + shIft88, MEST-OE + GANT61, MEST-OE + LY294002* groups. (d) qPCR showed relative *Gli2* and *Mest* mRNA expression in TSPCs under different treatment during in vitro osteogenic or chondrogenic differentiation. Data are presented as means ± SD of three independent assays. Statistical analyses were performed by one-way ANOVA analyses with Tukey's post hoc test for multiple comparisons. ***P < 0.005; ****P < 0.001.

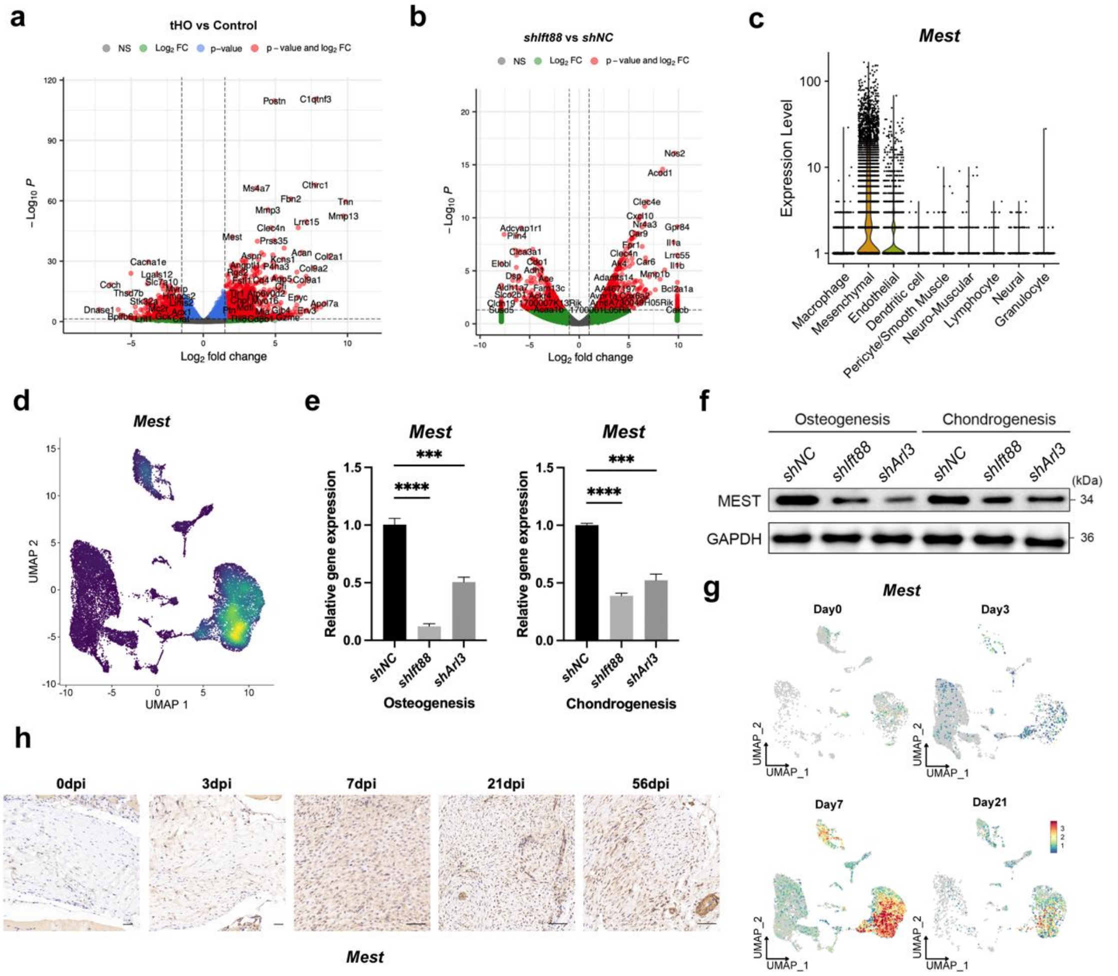


**Figure S4.** (a) The volcano plot visualizes the differential expressed genes in injured site of the tHO mice 3 weeks after burn/tenotomy compared to the uninjured control mice. (b) RNA-seq performed between *shIft88* and *shNC* TSPCs after 3-7 days of in vitro osteogenic differentiation and differential expressed genes are visualized. (c) The expression levels of *Mest* in each cluster are shown by violin plot. (d) The expression levels of *Mest* in each cluster are shown by UMAP plot. (e) qPCR shows relative *Mest* mRNA expression in TSPCs with or without *shIFT88* or *shARL3* treatment during in vitro osteogenic (left) or chondrogenic (right) differentiation. (f) Western blot shows MEST protein expression levels in TSPCs with or without *shIFT88* or *shARL3* treatment during in vitro osteogenic or chondrogenic differentiation. (g) UMAP plot shows the expression of *Mest* at different time points (Day 0, 3, 7, 21) during tHO development. (h) Immunohistochemistry images demonstrates MEST in tHO tendon tissues at different time post burn/tenotomy injury (scale bar 100 μm). Data are presented as means ± SD of three independent assays. Statistical analyses were performed by one-way ANOVA analyses with Tukey's post hoc test for multiple comparisons. ***P < 0.005; ****P < 0.001.

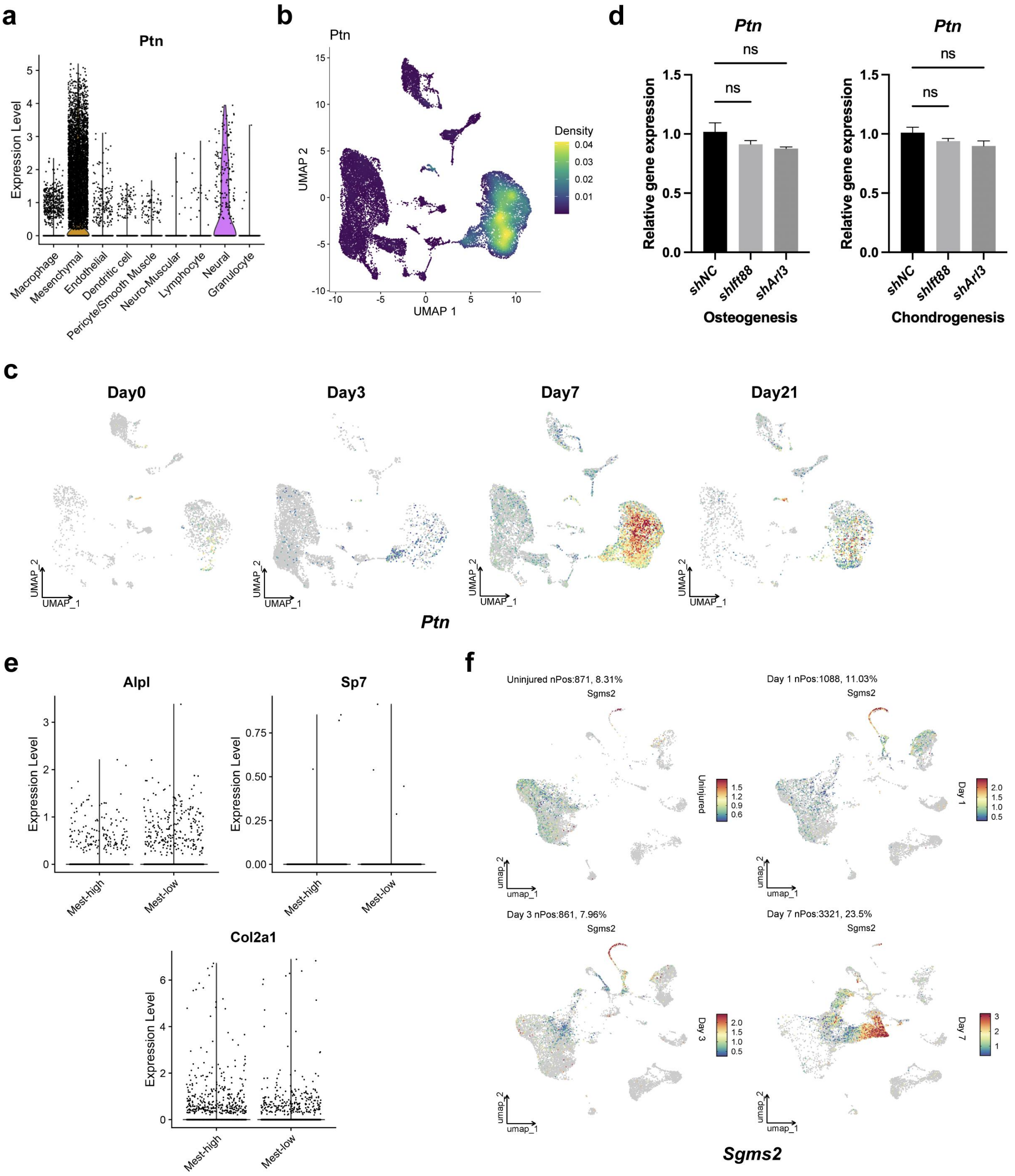


**Figure S5.** (a) The expression level of *Ptn* in each cluster are shown by violin plot. (b) The expression density of *Ptn* in each cluster are shown by UMAP plot. (c) UMAP plot shows the expression of *Ptn* in different time points (Day 0 ,3, 7, 21) during tHO development. (d) PCR showed relative *Ptn* mRNA expression in TSPCs with or without *shIFT88* or *shARL3* treatment during in vitro osteogenic or chondrogenic differentiation. (e) The expression level of *Alpl, Sp7* and *Col2a1* in Mest-high and Mest-low subclusters were visualize by violin plots. (f) UMAP plot shows the expression of *Sgms2* in different cells from *Nse-Bmp4* mice injured muscle in different groups during tHO development. Statistical analyses were performed by one-way ANOVA analyses with Tukey's post hoc test for multiple comparisons. ns, P > 0.05.

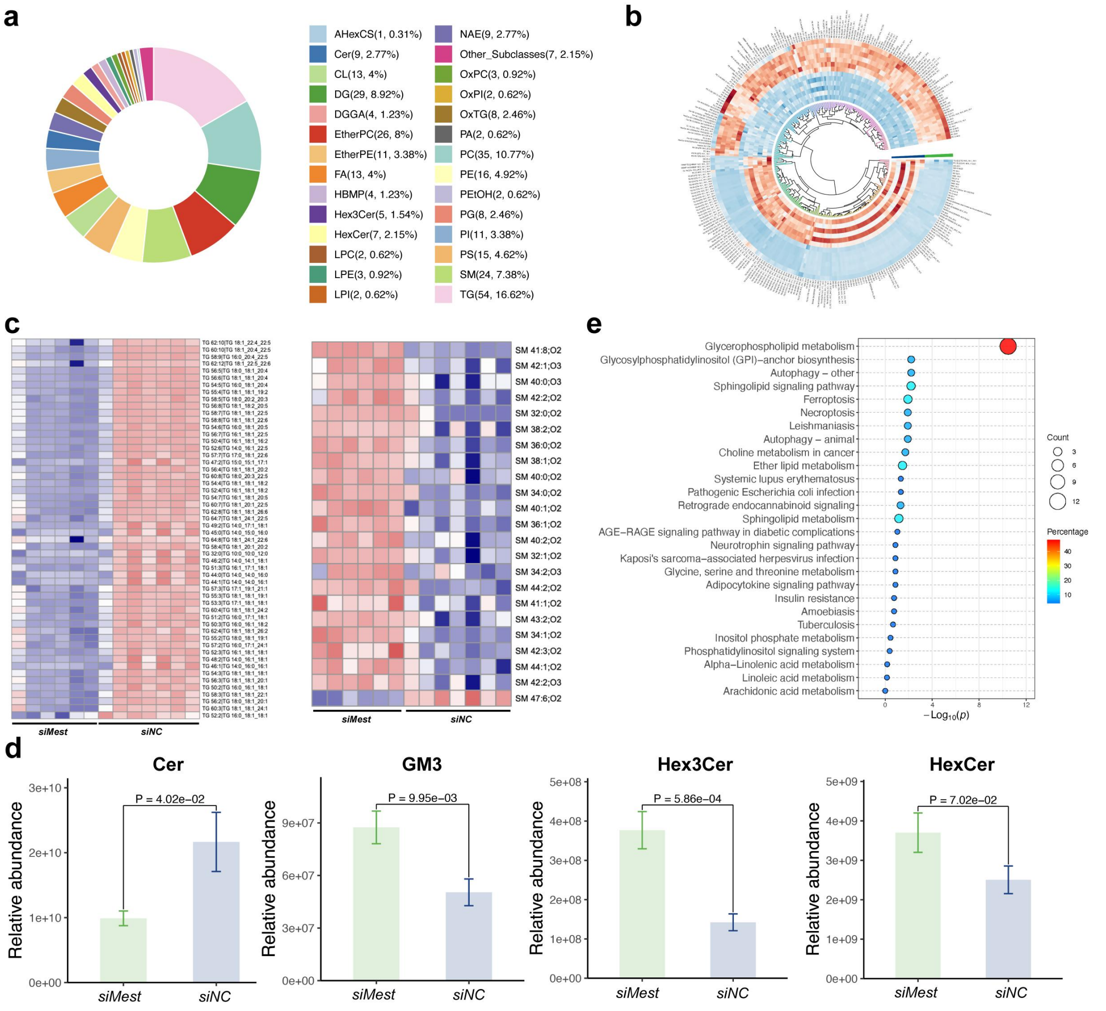


**Figure S6.** (a) Circular diagram of the subclass statistics of differential lipids. (b) Circular heat map of hierarchical clustering of significantly differential lipids between *siMest* and *siNC* groups. (c) The heatmap shows the different expression of 53 subclasses of triglycerides and 22 subclasses of sphingomyelins between *siMest* and *siNC* group. (d) Relative abundance of sphingolipids (Cer, GM3, Hex3Cer, and HexCer) are significantly different in *siMest* and *siNC* group. (e) KEGG enrichment analysis of differential expressed lipids in *siMest* group shows top 20 enriched pathways.

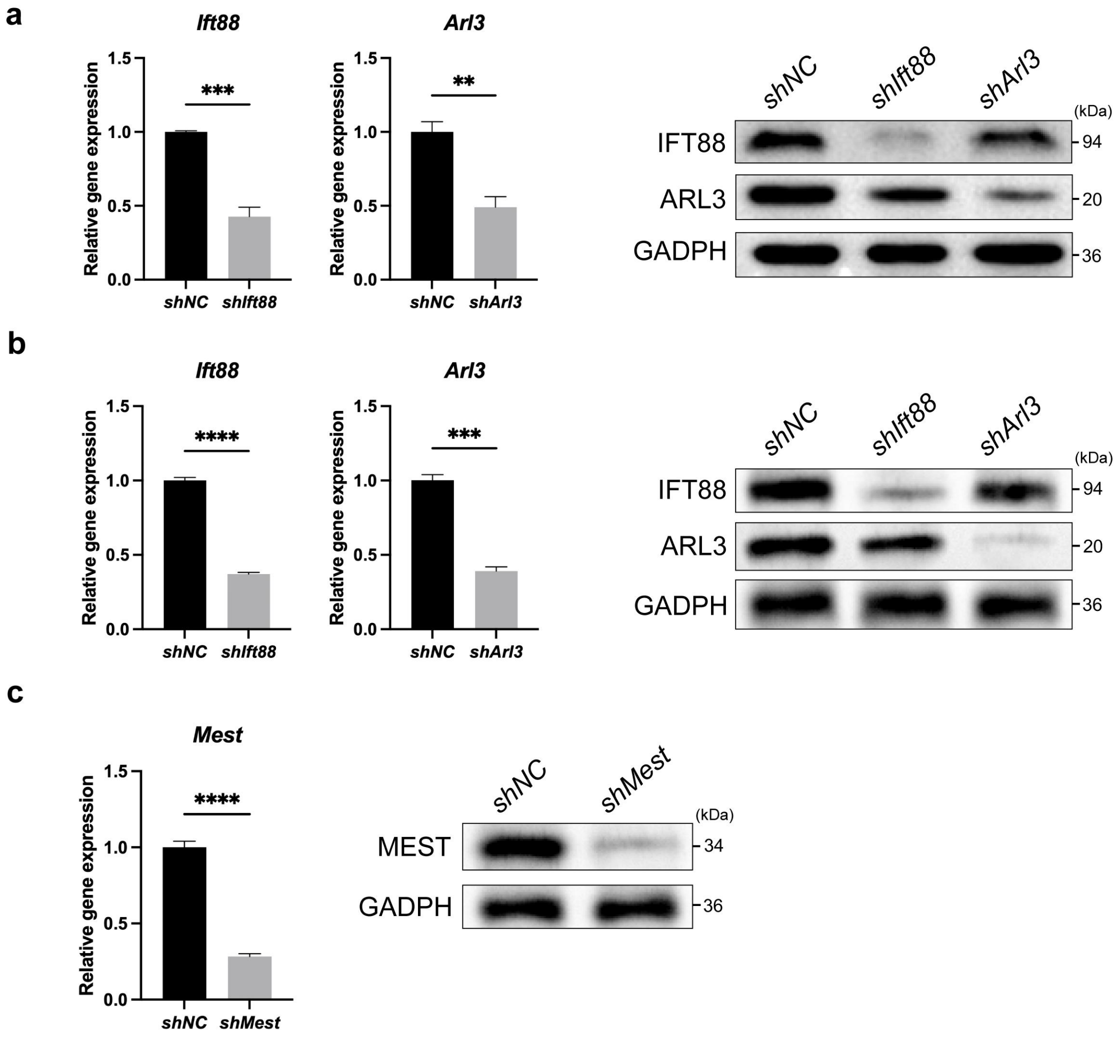


**Figure S7.** (a) Transfection efficiency of *shIft88* and *shArl3* in TSPCs was assessed by qPCR and Western blot. (b) Transfection efficiency of *shIft88* and *shArl3* in tHO injured site was assessed by qPCR and Western blot. (c) Transfection efficiency of *shMest* in tHO injured site was assessed by qPCR and Western blot. Data are presented as means ± SD of three independent assays. Statistical analyses were performed by Student's t-test for two-group comparison. $^{**}P < 0.01$; $^{***}P < 0.005$; $^{****}P < 0.001$.

**Table S1.** Genes upregulated in the mesenchymal stem cell subcluster at 7 and 21 days during tHO in mice (GSE126060) by scRNA-seq; genes upregulated in injured tissue at 21 days of tHO mice (GSE233201) by bulk RNA-seq; and genes downregulated in TSPCs with ciliary gene *Ift88* knockdown by bulk RNA-seq.

**Table S2.** The 316 differential expressed lipids between *siMest* group and *siNC* group.

**Table S3**. List of antibodies used in this study.

| **Antibody** | **Company** | **Rate** |
|---|---|---|
| ARL13B | #17711-1-AP, Proteintech | 1:500 (IF) |
| Acetyl-α-Tubulin (Lys40) | #12152, CST | 1:800 (IF) |
| PDGFRα | #sc-338, Santa Cruz | 1:100 (IF) |
| CD31 | #11265-1-AP, Proteintech | 1:500 (IF) |
| F4/80 | #28463-1-AP, Proteintech | 1:500 (IF) |
| TNMD | #PA5-112767, Invitrogen | 1:100 (IF) |
| α-SMA | #14395-1-AP, Proteintech | 1:500 (IF) |
| OCN | #23418-1-AP, Proteintech | 1:400 (IF) |
| COL2A1 | #28459-1-AP, Proteintech | 1:200 (IF) |
| MEST | #11118-1-AP, Proteintech | 1:250 (IF); 1:200 (IHC); 1:1000 (WB) |
| GLI2 | #28245-1-AP, Proteintech | 1:200 (IHC); 1:2000 (WB) |
| | #PA5-47500, Invitrogen | 1 µg/10^6 cells (ChIP) |
| PTCH1 | #ab53715, abcam | 1:100 (IHC); 1:1000 (WB) |
| SMO | #20787-1-AP, Proteintech | 1:1000 (IHC) |
| SHH | #20697-1-AP, Proteintech | 1:2000 (WB) |
| IFT88 | #13967-1-AP, Proteintech | 1:6000 (WB) |
| ARL3 | #10961-1-AP, Proteintech | 1:1000 (WB) |
| PI3K | #ab191606, abcam | 1:1000 (WB) |
| p-PI3K | #ab278545, abcam | 1:2000 (WB) |
| AKT | #ab8805, abcam | 1:2000 (WB) |
| p-AKT | #ab38449, abcam | 1:1000 (WB) |
| GAPDH | #10494-1-AP, Proteintech | 1:10000 (WB) |

| | | |
|---|---|---|
| β-Actin | #20536-1-AP, Proteintech | 1:6000 (WB) |
| SGMS2 | #PA5-113988, Invitrogen | 1:1000 (WB) |

**Table S4**. List of shRNA and siRNA used in this study.

| **Gene** | **Sequence (5’-3’)** |
|---|---|
| *shARL3* | AAGCTGAATGTATGGGACATT |
| *shIFT88* | GCCCTCAGATAGAAAGACCAA |
| *shMest* | CACCGGCCATTGGATCCTAT |
| *shNC* | CCTAAGGTTAAGTCGCCCTCG |
| *siMest-1* | UGGUCAUCCAGAAUCGACACUGUGG |
| *siMest-2* | GAGGAUCCCAUGGGCUUCUUGAAUG |
| *siGli2* | GGAAAACUUCAACAAUACATT |
| *siNC* | UGGGUUCUACAGACGCUACACAUGG |

**Table S5**. List of primers used in this study for qPCR and ChIP-qPCR.

| **Gene** | **Primer** | **Sequence (5’-3’)** |
|---|---|---|
| *Gapdh* | Forward | ACTCCACTCACGGCAAATTC |
| | Reverse | TCTCCATGGTGGTGAAGACA |
| *Shh* | Forward | CTGGCTCGCCTGGCTGTG |
| | Reverse | CCACGGAGTTCTCTGCTTTCAC |
| *Ptch1* | Forward | GGCTGGCTGGCGTCCTG |
| | Reverse | CATCATCCACACCAACACCAAGAG |
| *Alpl* | Forward | CACGGCGTCCATGAGCAGAAC |
| | Reverse | CAGGCACAGTGGTCAAGGTTGG |
| *Runx2* | Forward | TCCCAGGCAGGCACAGTCTTC |
| | Reverse | AGCGGCGTGGTGGAGTGG |
| *Bglap* | Forward | AGGGCAGCGAGGTAGTGAAGAG |
| | Reverse | GCCGATGTGGTCAGCCAACTC |
| *Sp7* | Forward | CCTCTGCGGGACTCAACAAC |
| | Reverse | AGCCCATTAGTGCTTGTAAAGG |
| *Col2a1* | Forward | GGTCCTCCTGGTCCTGGCATC |
| | Reverse | CGTGCTGTCTCAAGGTACTGTCTG |

| | | |
|---|---|---|
| *Sox9* | Forward | ATGACCGACGAGCAGGAGAAGG |
| | Reverse | CCGAGGGACAGGGCGAACC |
| *Acan* | Forward | GGAGACCCAGACAGCAGAAACAAC |
| | Reverse | GCAGGTGGCTCCATTCAGACAAG |
| *Ift88* | Forward | TTCAGCAAGCAGTGAGAACCAGTC |
| | Reverse | TGTCCCTGTCATCGGTCTTCCC |
| *Arl3* | Forward | TGTGTGCCCGTGCTCATCTTTG |
| | Reverse | TGGACGCCCTCGCCTGTG |
| *Gli2* | Forward | GCTCCACACACCCGCAACAC |
| | Reverse | CTCAGCATCGTCACTTCGGTCAG |
| *Mest* (qPCR) | Forward | CTTAGGCTTTGGCTTCAGTGACAA |
| | Reverse | AGGTTGATTCTGCGGTTCTGGAG |
| *Mest* (ChIP-qPCR) site 1 | Forward | CCGCATTACGTGTTCTGATGA |
| | Reverse | GGGATCTAACCTGTGGGTATCT |
| *Mest* (ChIP-qPCR) site 2 | Forward | TATGTCTTCCAGGGTCTAGGG |
| | Reverse | GGATGACAAACGTCTGAGAGAG |
